# Reconstruction of developmental landscapes by optimal-transport analysis of single-cell gene expression sheds light on cellular reprogramming

**DOI:** 10.1101/191056

**Authors:** Geoffrey Schiebinger, Jian Shu, Marcin Tabaka, Brian Cleary, Vidya Subramanian, Aryeh Solomon, Siyan Liu, Stacie Lin, Peter Berube, Lia Lee, Jenny Chen, Justin Brumbaugh, Philippe Rigollet, Konrad Hochedlinger, Rudolf Jaenisch, Aviv Regev, Eric S. Lander

**Affiliations:** Broad Institute of MIT and Harvard, Cambridge, MA 02142, USA; Whitehead Institute for Biomedical Research, Cambridge, MA 02142, USA; Computational and Systems Biology Program, MIT, Cambridge, MA 02142, USA; Harvard-MIT Division of Health Sciences and Technology, Cambridge, MA 02139 USA; Cancer Center, Massachusetts General Hospital, Boston, MA 02114 USA; Department of Biology, Massachusetts Institute of Technology, Cambridge, MA 02139, USA; Department of Molecular Biology, Center for Regenerative Medicine and Cancer Center, Massachusetts General Hospital, Boston, MA 02114, USA; Department of Stem Cell and Regenerative Biology, Harvard University, Cambridge, MA 02138, USA; Harvard Stem Cell Institute, Cambridge, MA 02138, USA; Harvard Medical School, Boston, MA 02115, USA; MIT Center for Statistics, Massachusetts Institute of Technology, Cambridge, MA 02139, USA; Department of Mathematics, Massachusetts Institute of Technology, Cambridge, MA 02139, USA; Howard Hughes Medical Institute, Chevy Chase, MD, USA; Department of Systems Biology Harvard Medical School, Boston, MA 02125, USA; Biochemistry Program, Wellesley College, Wellesley, 02481, MA, USA

## Abstract

Understanding the molecular programs that guide cellular differentiation during development is a major goal of modern biology. Here, we introduce an approach, WADDINGTON-OT, based on the mathematics of optimal transport, for inferring developmental landscapes, probabilistic cellular fates and dynamic trajectories from large-scale single-cell RNA-seq (scRNA-seq) data collected along a time course. We demonstrate the power of WADDINGTON-OT by applying the approach to study 65,781 scRNA-seq profiles collected at 10 time points over 16 days during reprogramming of fibroblasts to iPSCs. We construct a high-resolution map of reprogramming that rediscovers known features; uncovers new alternative cell fates including neural- and placental-like cells; predicts the origin and fate of any cell class; highlights senescent-like cells that may support reprogramming through paracrine signaling; and implicates regulatory models in particular trajectories. Of these findings, we highlight *Obox6*, which we experimentally show enhances reprogramming efficiency. Our approach provides a general framework for investigating cellular differentiation.

## Introduction

In the mid-20^th^ century, Waddington introduced two images to describe cellular differentiation during development: first, trains moving along branching railroad tracks and, later, marbles following probabilistic trajectories as they roll through a developmental landscape of ridges and valleys (**Figure 1A,B**) [*1, 2*]. These metaphors have powerfully shaped biological thinking in the ensuing decades. The recent advent of massively parallel single-cell RNA sequencing (scRNASeq) [*3-7*] now offers the prospect of empirically reconstructing and studying the actual “landscapes”, “fates” and “trajectories” associated with complex processes of cellular differentiation and de-differentiation—such as organismal development, long-term physiological responses, and induced reprogramming—based on snapshots of expression profiles from heterogeneous cell populations undergoing dynamic transitions [*6-11*].

**Figure 1.**
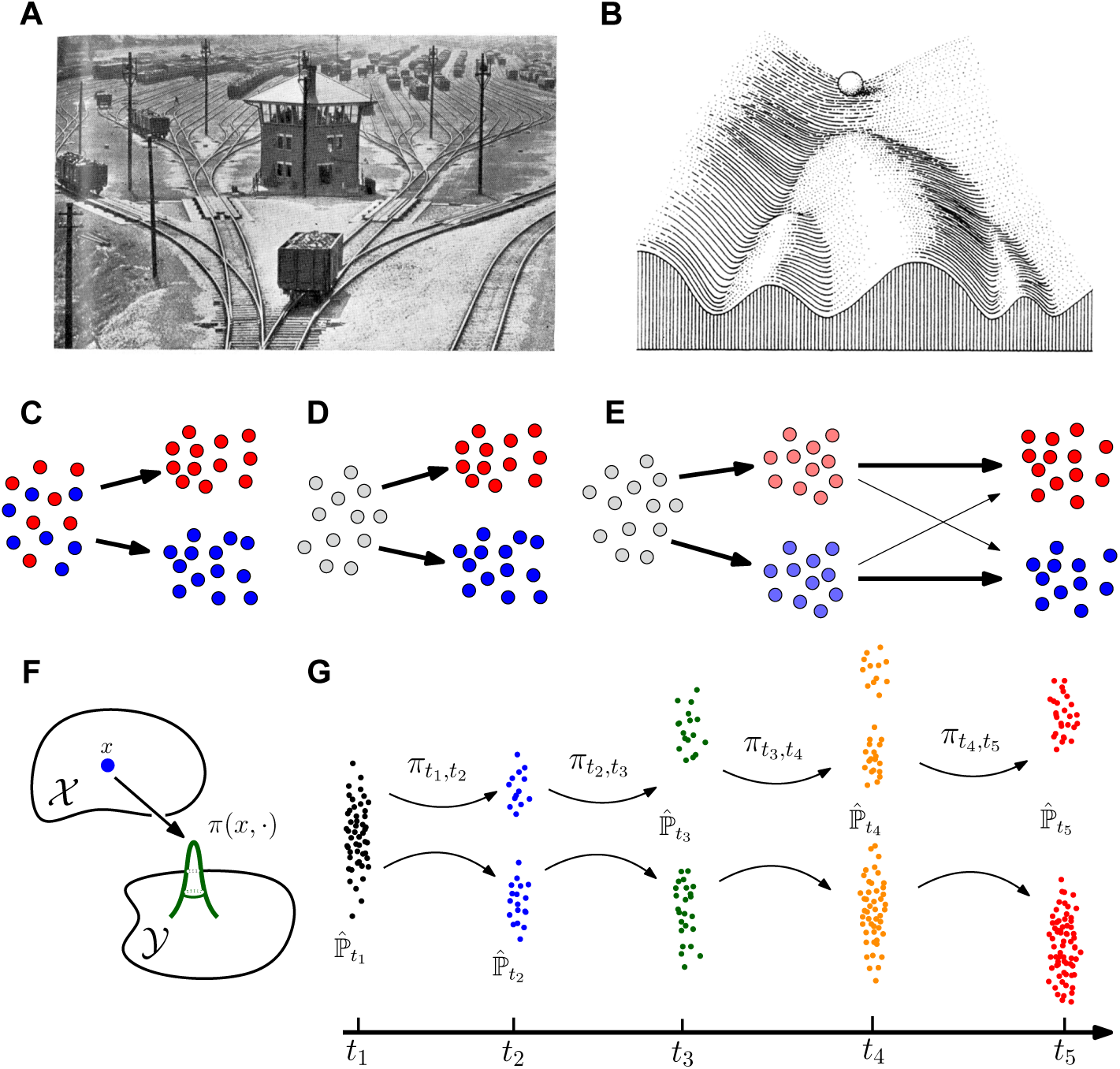
Schematic representations of cellular differentiation. (**A-B**) Waddington’s classic analogies of cells undergoing differentiation, initially (1936) illustrated by railroad cars on switching tracks (**A**) and later (1957) by marbles rolling in a landscape (**B**), with trajectories shaped by hills and valleys. (**C-E**) Differentiation processes in which the ultimate fate of individual cells (filled dots) is (**C**) predetermined (**D**) not predetermined, or (**E**) progressively determined. Arrows indicate possible transitions, and color represents cell fate, with red and blue indicating distinct fates, light red and light blue indicating partially determined fates, and grey indicating undetermined fate. (**F**) Illustration of transported mass. A transport map, π, describes how a point x at one stage (X) is redistributed across all points (denoted by “·”) at the subsequent stage (Y). (**G**) Transport maps computed from a time series of samples taken from a time-varying distribution. Between each pair of time points, a transport map 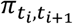 redistributes the cells observed at time *t_i_* to match the distribution of cells 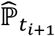 observed at time *t_i_*_+1_.

To understand such processes in detail, we need general approaches to answer key questions. For any given system, we would like to know: What classes of cells are present at each stage? For the cells in each class, what was their origin at earlier stages, what are their potential fates at later stages, and what is the actual outcome of a given cell? To what extent are events along a path synchronous or asynchronous? What are the genetic regulatory programs that control each path? What are the intercellular interactions between classes of cells? Answering these questions would provide insights into the nature of developmental processes (**Figure 1C-E**): How deterministic or stochastic is the process—that is: if, and how early, does it become determined that a particular cell or an entire cell class is destined to a specific fate? For a given origin and target fate, is there only a single path to the target, or are there multiple developmental paths? To what extent is the process cell-intrinsic, driven by intracellular mechanisms that do not require ongoing external inputs, or externally regulated, being affected by other contemporaneous cells? For artificial processes such as induced reprogramming, there are additional questions: What off-target cell classes arise? To what extent do cells activate normal developmental programs vs. unnatural hybrid programs? How can the efficiency of reprogramming be improved?

Experimental approaches to such questions have typically involved studying bulk populations or identifying subsets of cells based on activation of one or a few genes at a specific time (*e.g.*, reporter genes or cell-surface markers) and tracing their subsequent fate. These experiments are severely limited, however, by the need to choose subsets of cells *a priori* and develop distinct reagents to study each subset. For example, studies of cellular reprogramming from fibroblasts to induced pluripotent cells (iPSCs) have largely relied on RNAand chromatinprofiling studies of bulk cell populations, together with fate-tracing of cells based on a limited set of markers (e.g., *Thy1* and *CD44* as markers of the fibroblast state, and *ICAM1*, *Oct4*, and *Nanog* as markers of partial reprogramming) [*12-16*].

Computational approaches based on single-cell gene expression profiles offer a complementary approach with broader molecular scope, because one can readily define classes of cells based on any expression profile at any stage. The remaining challenge is to reliably infer their trajectories across stages.

Several pioneering papers have introduced methods to infer cellular trajectories [*9, 10, 17-29*]. Early studies recognized that cellular profiles from heterogeneous populations can provide information about the temporal order of asynchronous processes—enabling intermediate transitional cells to be ordered in “pseudotime” along “trajectories”, based on their state of cell differentiation [*18*]. Some approaches relied on *k*-nearest neighbor graphs [*18*] or binary trees [*9*]. More recently, diffusion maps have been used to order cell state transitions. In this case, single-cell profiles are assigned to densely populated paths through diffusion map space [*20, 21*]. Each such path is interpreted as a transition between cellular fates, with trajectories determined by curve fitting, and cells are “pseudotemporally ordered” based on the diffusion distance to the endpoints of each path. Whereas initial efforts focused mostly on single paths, more recent work has grappled with challenges of branching, which is critical for understanding developmental decisions [*10, 11, 21*].

While these pioneering approaches have shed important light on various biological systems, many important challenges remain. First, because many methods were initially designed to extract information about stationary processes (such as the cell cycle or adult stem cell differentiation) in which all stages exist simultaneously, they neither directly model nor explicitly leverage the temporal information in a developmental time course [*29*]. Second, a single cell can undergo multiple temporal processes at once. These processes can dramatically impact the performance of these models, with a notable example being the impact of cell proliferation and death [*29*]. Third, many of the methods impose strong structural constraints on the model, such as one-dimensional trajectories and zero-dimensional branch points. This is of particular concern if development follows the flexible “marble” rather than the regimented “tracks” models, in Waddington’s frameworks (**Figure 1A,B**).

Here, we describe a conceptual framework intended to reflect Waddington’s image of marbles rolling within a development landscape. It aims to capture the notion that cells at any position in the landscape have a *distribution* of both probable origins and probable fates. It seeks to reconstruct both the landscape and probabilistic trajectories from scRNA-seq data at various points along a time course. Specifically, it uses time-course data to infer how the probability distribution of cells in gene-expression space evolves over time, by using the mathematical approach of Optimal Transport (OT). We implement this framework in a method, Waddington-OT, and demonstrate its capabilities in the context of reprogramming of fibroblasts to induced pluripotent stem cells (iPSCs). Waddington-OT readily rediscovers known biological features of reprogramming, including that successfully reprogrammed cells exhibit an early loss of fibroblast identity, maintain high levels of proliferation, and undergo a mesenchymal-to-epithelial transition before adopting an iPSC-like state [*12*]. In addition, by exploiting single-cell resolution and the new model, it also extends these results by (1) identifying alternative cell fates, including senescence, apoptosis, neural identity, and placental identity; (2) quantifying the portion of cells in each state at each time point; (3) inferring the probable origin(s) and fate(s) of each cell and cell class at each time point; (4) identifying early molecular markers associated with eventual fates; and (5) using trajectory information to identify transcription factors (TFs) associated with the activation of different expression programs. In particular, we propose TFs that are putative regulators of neural identity, placental identity, and pluripotency during reprogramming, and we experimentally demonstrate that one such TF, *Obox6*, enhances reprogramming efficiency. Together, the data provide a high-resolution resource for studying the roadmap of reprogramming, and the methods provide a general approach for studying cellular differentiation in natural or induced settings.

## Results

### Reconstruction of Probabilistic Trajectories by Optimal Transport

To develop a probabilistic framework for reconstructing developmental trajectories from scRNA-Seq data along a time course, we consider each cell at each time *t* to be drawn from a time-dependent probability distribution *P*_*t*_ in a high-dimensional gene-expression space. By sampling *P*_*t*_ at various time points, we wish to infer how *P*_*t*_ evolves over time. We imagine *P*_*t*_ as being reshaped by “transporting” mass from one time to the next (**Figure 1F,G**). In practice, we aim to redistribute the masses of the specific cells sampled at time *t* to form the masses of the cells sampled at time *t+1*. Although the cells in the latter set are not the literal descendants of those in the former set (because the sampling process is destructive), we expect that they are *representative* of the descendants, provided enough cells are sampled. Moreover, if the gap between samples is short enough, we expect the distributions will tend to flow continuously through space so that cells at time *t* tend to have their mass redistributed to cells at time *t+1* with similar gene expression patterns (rather than most cells remaining stationary and a minority making huge leaps). To reconstruct the time-dependent evolution of *P*_*t*_, we thus seek to find couplings *π*_*t,t+1*_ (called transport maps) that redistribute mass between the sampled distributions that minimize the “transport cost” between two samples distributions—defined in terms of the sum across all cells of mass x squared distance moved, for an appropriate distance metric in gene expression space (**Figure 1F,G**).

This problem belongs to the classical field of “optimal transport,” originally developed by Monge in the 1780s to redistribute piles of dirt to build fortifications with minimal work and soon applied by Napoleon in his campaign in Egypt [*30*]. Large-scale optimal transport problems have traditionally been difficult to solve, but a recently invented class of algorithms called entropically regularized transport [*31, 32*] dramatically accelerates numerical computations and helps prevent overfitting. Our problem differs from classical optimal transport in one important respect: unlike dirt, cells can proliferate. We therefore allow the mass at each point to change over time, based on a gene-expression signature of cell proliferation (*Appendix S1*).

If we assume that a cell’s fate depends only on its current position (that is, is memoryless), we can infer couplings between more distant time points by composing transport maps between the intermediate time points. In this case, differentiation follows a velocity field on gene expression space, and the potential inducing this velocity field corresponds directly with Waddington’s landscape.

We define trajectories in terms of “descendant distributions” and “ancestor distributions” as follows. For any set S of cells at time *t*_*1*_, its “descendant distribution” at a later time *t*_*2*_ refers to the *mass distribution* over all cells at time *t*_*2*_ obtained by transporting S according to the transport maps. Branching events, for example, are revealed by the emergence of bimodality in the descendant distribution. The “ancestor distribution” at an earlier time *t*_*0*_ is defined as a mass distribution over all cells at time *t*_*0*_, obtained by transporting S in the opposite direction (that is, as though one “rewinds” time). The “trajectory” *from* S refers to the sequence of descendant distributions at each subsequent time point, and the trajectory *to* S similarly refers to the sequence of ancestor distributions. For expositional convenience below, we will sometimes refer simply to the “ancestors” (and, synonymously, “origin”), “descendants” (and “fate”), and trajectories of cells. These terms will always refer to a *distribution* over a set of expression patterns corresponding to observed cells that serve as *proxies* for the actual ancestors or descendants.

In summary, for any cell or set of cells, the optimal-transport analysis provides a distribution over representative ancestors and descendants at any other time. This information allows us to readily assess whether any set of cells is “biased” toward a given outcome—that is, whether its descendants are enriched for the outcome (compared to another set of contemporaneous cells).

### Implementation of optimal transport in WADDINGTON-OT

We implemented this theoretical framework as a method, called WADDINGTON-OT, for exploratory analysis of developmental landscapes and trajectories (*Appendix S2*). We are preparing a software package incorporating the methods for public distribution. The method includes:

(1) Performing optimal-transport analyses on scRNA-seq data from a time course, by calculating optimal-transport maps and using them to find ancestors, descendants and trajectories for any set S of cells.

(2) Learning regulatory models from optimal transport, based on a global model (built by sampling pairs of cells at time *t* and *t+1* according to their transport probabilities and using the expression levels of TFs at the earlier time point to predict non-TF expression at the later time point) and local enrichment analysis (identifying TFs enriched in cells having many *vs*. few descendants (>80% *vs*. <20% mass) in a target cell population).

(3) Defining gene modules, by partitioning genes based on correlated expression across cells, and cell clusters, by partitioning cells based on graph clustering [*33, 34*] (following dimensionality reduction by diffusion maps [*20, 35*].

(4) Visualizing the developmental landscape (including the cells, ancestors, descendants, trajectories, gene modules and cell clusters) by representing the high-dimensional geneexpression data in two dimensions (using Force-Directed Layout Embedding (FLE)) [*36-38*].

### Reprogramming to iPSCs as a test case for analysis of developmental landscapes

To test these ideas, we applied WADDINGTON-OT to reprogramming of fibroblasts to iPSCs [*39-42*]. This system provides ample opportunity to compare results with previous studies, while also exploring open questions.

Most of what is known about reprogramming from fibroblasts to iPSCs comes from studies of bulk cell populations. A few recent studies have applied scRNA-Seq, but they have involved only several dozen cells or several dozen genes [*13, 43*]. Studies have proposed that reprogramming involves two “transcriptional waves,” with gain of proliferation and loss of fibroblast identity followed by transient activation of developmental regulators and gradual activation of embryonic stem cell (ESC) genes [*12*]. Some studies [*16,44,45*], including from our own group [*45*], have noted strong upregulation of lineage-specific genes from unrelated lineages (*e.g.*, related to neurons), but it has been unclear whether this largely reflects disorganized gene activation by TFs or coherent differentiation of specific (off-target) cell types [*45*]. We sought to explore how much could be learned from probabilistic analysis of a largescale scRNA-seq of the developmental landscape.

We collected scRNA-seq profiles of 65,781 cells across a 16-day time course of iPSC induction, under two conditions (**Figure 2A,B**) (*Appendix S3*). Because the derivation of iPSCs by primary infection is very inefficient, we used a more efficient “secondary” reprogramming system [*46*].We obtained mouse embryonic fibroblasts (MEFs) from a single female embryo homozygous for ROSA26-M2rtTA, which constitutively expresses a reverse transactivator controlled by doxycycline (Dox), a Dox-inducible polycistronic cassette carrying *Pou5f1* (*Oct4*), *Klf4*, *Sox2*, and *Myc (OKSM)*, and an EGFP reporter incorporated into the endogenous *Oct4* locus (Oct4IRES-EGFP). We plated MEFs in serum-containing induction medium, with Dox added on day 0 to induce the OKSM cassette (Phase-1(Dox)). Following Dox withdrawal at day 8, we transferred cells to either serum-free N2B27 2i medium (Phase-2(2i)) or maintained the cells in serum (Phase-2(serum)). Oct4-EGFP^+^ cells emerged on day 10 as a reporter for “successful” reprogramming to endogenous Oct4 expression (**Figure 2C**). We collected single or duplicate samples at the various time points (**Figure 2A**), generated single cell suspensions and performed scRNA-Seq (**table S1**, **Figure S1**). We also collected samples from established iPSC lines reprogrammed from the same MEFs, maintained in either 2i or serum conditions.Overall, we profiled 68,339 cells to an average depth of 38,462 reads per cell (**table S1**). After discarding cells with less than 1,000 genes detected, we retained a total of 65,781 cells, with a median of 2,398 genes and 7,387 unique transcripts per cell.

**Figure 2.**
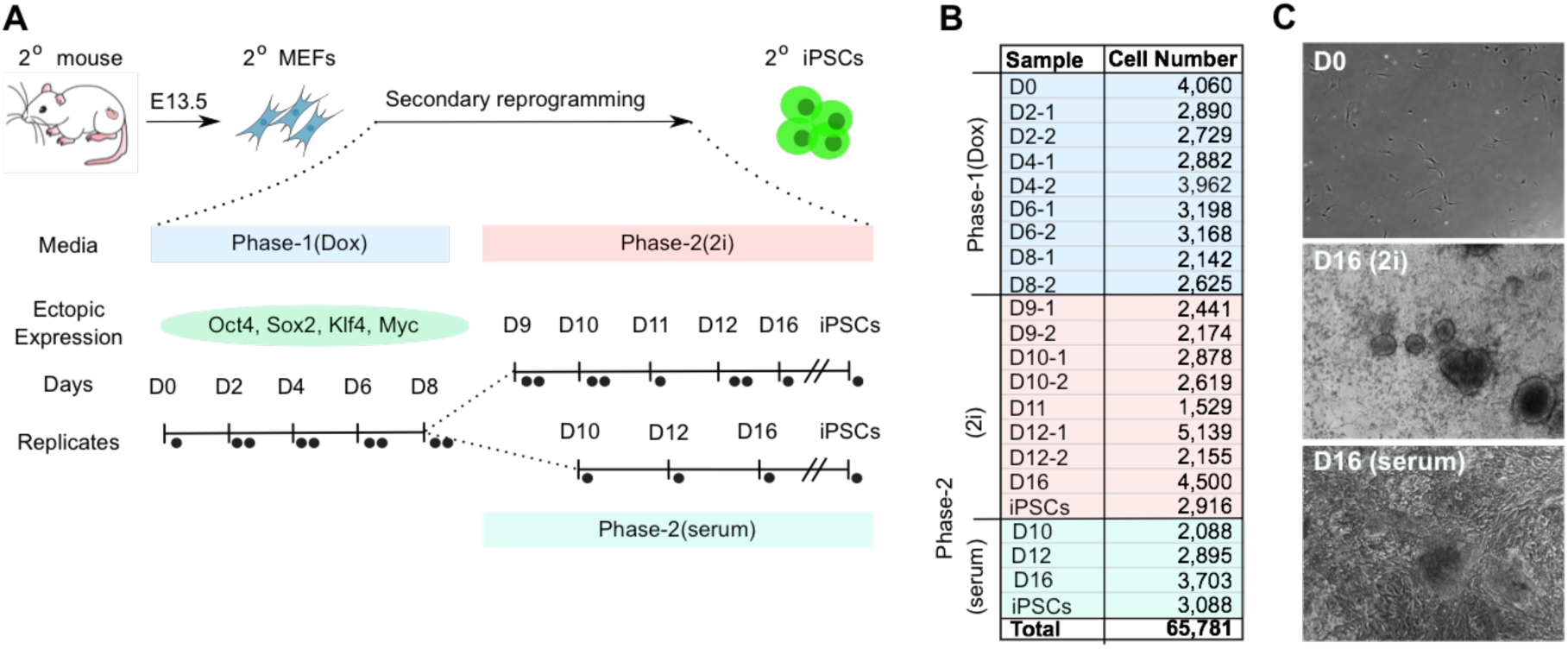
Experimental design of scRNA-Seq time course during iPSC reprogramming. (**A**) Representation of reprogramming procedure and time points of sample collection. (Top) Mouse embryos (E13.5) were dissected to obtain secondary MEFs (2^o^ MEF), which were reprogrammed into iPSCs. In Phase-1 of reprogramming (light blue; days 0-8), doxycycline (Dox) was added to the media to induce ectopic expression of reprogramming factors (*Oct4*, *Klf4*, *Sox2*, and *Myc*). In Phase-2 (days 9-16), Dox was withdrawn from the media, and cells were grown either in the presence of 2i (light red) or serum (light green). Samples were also collected from established iPSC lines reprogrammed from the same 2^o^ MEFs, maintained in either 2i or serum conditions (far right in each time course). Individual dots along the time course indicate time points of scRNA-Seq collection, with two dots indicating biological replicates. (**B**) Number of scRNA-Seq profiles from each sample collection that passed quality control filters. (**C**) Bright field images of day 0 (Phase1-(Dox)) and day 16 cells during reprogramming in (Phase-2(2i)) and (Phase-2(serum)) culture conditions.

### The reprogramming landscape reveals relationships among biological features

Using WADDINGTON-OT, we generated a transport map across the cells in the time course. Based on similarity of expression profiles, we partitioned the 16,339 detected genes into 44 gene modules and the 65,781 cells into 33 cell clusters. Some of the clusters contain cells from more than one time point, reflecting asynchrony in the reprogramming process.

We explored the landscape of reprogramming by identifying cell subsets of interest (*e.g.*, successfully reprogrammed cells from day 16, or each of the cell clusters), studying the trajectories to and from these subsets (*e.g.*, characterizing the pattern of gene expression in ancestors at day 8 of successfully reprogrammed target cells at day 16), and considering contemporaneous interactions between them. We visualized the analyses in a two-dimensional embedding using FLE (**Figure 3A**), which we annotated in various ways (*Appendix S2*). FLE reflects better global structures in our data than other modes of visualization **(Figure S2)**.These annotations include time points and growth conditions (**Figure 3B,C**), gene modules (**Figure S3, S4, table S2**), cell clusters (**Figure 3D, Figure S4, table S3**), expression of gene signatures (curated gene sets associated with specific cell types, pathways, and responses, such as MEF identity, proliferation, pluripotency, and apoptosis; **Figure 3E, table S4**), expression of individual genes (**Figure 3F, Figure S5**), and ancestor and descendant distributions (**Figure 4**). Extensive sensitivity analysis showed that key biological results for the reprogramming data were largely robust to the details of the formulation. Finally, we compared the WADDINGTON-OT landscape to the landscapes produced by various graph-based methods (*Appendix S4*).

**Figure 3.**
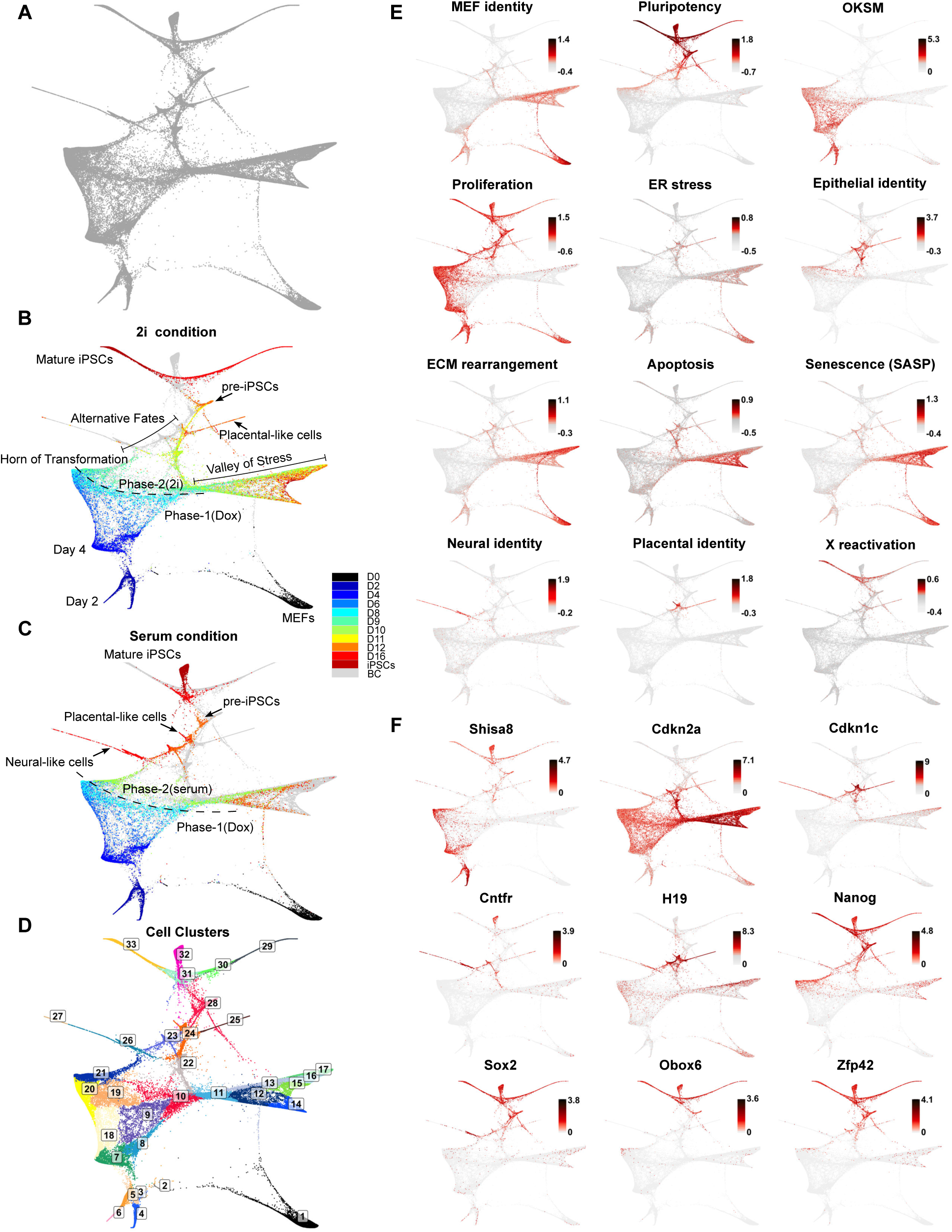
scRNA-Seq reprogramming landscape annotated in force-directed layout embedding (FLE). scRNA-Seq profiles of all 65,781 cells were embedded in two-dimensional space using FLE, and annotated with indicated features. (**A**) Unannotated layout of all cells. Each dot represents one cell. (**B,C**) Annotation by time point (color) and biological feature, with Phase-2 points from either (**B**) 2i condition or (**C**) serum condition. Phase-1 points appear in both (**B**) and (**C**).Individual cells are colored by day of collection, with grey points (BC, background color) representing Phase-2 cells from serum (in **B**) or 2i (in **C**). (**D**) Annotation by cell cluster. Cells were clustered on the basis of similarity in gene expression. Each cell is colored by cluster membership (with clusters numbered 1-33). (**E-F**) Annotation by gene signature (**E**) and individual gene expression levels (**F**). Individual cells are colored by gene signature scores (in **E**) or normalized expression levels (in **F**;, where E is the number of transcripts of a gene per 10,000 total transcripts). See Figure S3, 5 for additional genes and modules.

**Figure 4.**
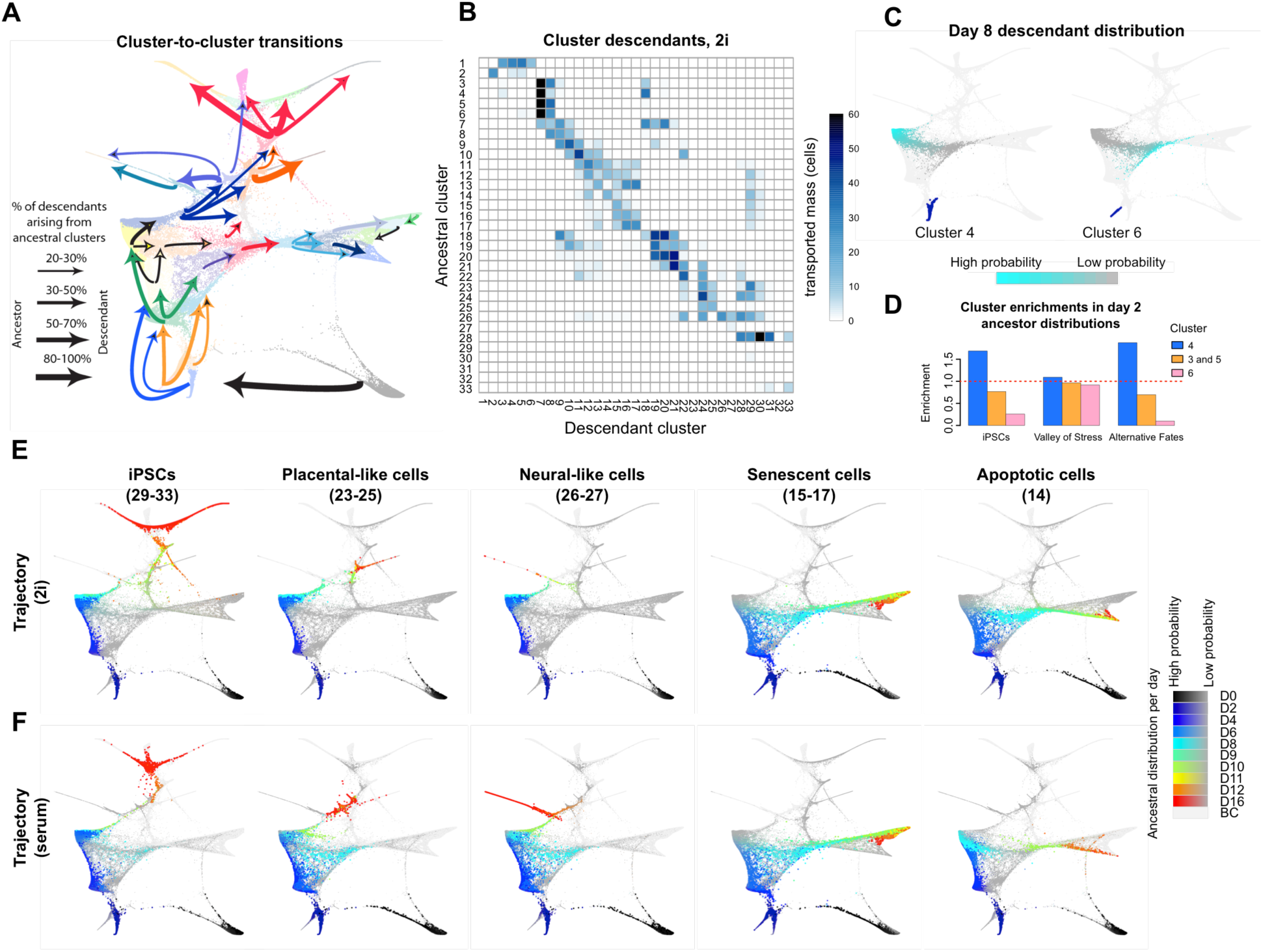
Ancestral origins and descendant fates inferred by optimal transport. **(A)** Schematic representation of the major cluster-to-cluster transitions (see Table S5 for details[BC17]). Individual arrows indicate transport from ancestral clusters to descendant clusters, with colors corresponding to the ancestral cluster. For each descendant cluster, arrows were drawn when at least 20% of the ancestral cells (at the previous time point) were contained within a given cluster (self-loops not shown). Arrow thickness indicates the proportion of ancestors arising from a given cluster. (**B**) Heatmap depiction of cluster descendants in 2i condition. In each row of the heatmap, color intensity indicates the number of descendant cells (“mass”, normalized to a starting population of 100 cells) transported to each cluster at the subsequent time point (see Table S5 for details). Clusters with highly-proliferative cells (*e.g.*, cluster 4) transport more total mass than clusters with lowly-proliferative cells (*e.g.*, cluster 14). (**C**) Depiction of divergent day 8 descendant distributions for two clusters of cells at day 2 (cluster 4 (left) and cluster 6 (right). Color intensity indicates the distribution of descendants at day 8, with bright teal indicating high probability fates and gray indicating low probability fates. (**D**) Enrichment of the ancestral distributions of iPSCs, Valley of Stress, and alternative fates (neuron-like and placenta-like) in clusters of day 2 cells. The red horizontal dashed line indicates a null-enrichment, where a cluster contributes to the ancestral distribution in proportion to its size. Cluster 4 has a net positive enrichment because its descendants are highly proliferative, while cluster 6 has a net negative enrichment because its descendants are lowly proliferative. (**E**) and (**F**) Ancestral trajectories of indicated populations of cells at day 16 (iPSCs, placental, neural-like cells, *etc*.) in serum (**E**) and 2i (**F**). Clusters used to define the indicated populations are shown in parentheses. Colors indicate time point.Sizes of points and intensity of colors indicate ancestral distribution probabilities by day (color bars, right; BC, background color, representing cells from the other culture condition).

We start with a brief overview of the landscape. Cell trajectories start at the lower right corner at day 0, proceed leftward to day 2 and then upward towards two regions that we dub the Valley of Stress and the Horn of Transformation (**Figure 3B**, **Figure 4A**). The Valley is characterized by signatures of cellular stress, senescence, and, in some regions, apoptosis (**Figure 3E**); it appears to be a terminal destination. By contrast, the Horn is characterized by increased proliferation, loss of fibroblast identity, a mesenchymal-to-epithelial transition (**Figure 3E**), and early appearance of certain pluripotency markers (*e.g.*, *Nanog* and *Zfp42*, **Figure 3F**), which are predictive features of successful reprogramming [*47*]. Some of the cells in the Horn proceed toward pre-iPSCs by day 12 and iPSCs by day 16, while others encounter alternative fates of placental-like development and neurogenesis (in serum, but not 2i condition; **Figure 3B,C**). We next present a more detailed account of the landscape.

### Predictive markers of reprogramming success are detectable by day 2

The vast majority (>98%) of cells at day 0 fall into a single cluster characterized by a strong signature of MEF identity, with clear bimodality in the proliferation signature (**Figure S6A**). By day 2 after Dox treatment, cells show high levels of expression of the OKSM cassette and have begun to diverge in their responses (clusters 3, 4, 5, 6, **Figure 3D**). Overall, they score highly for expression signatures of proliferation, MEF identity, and endoplasmic reticulum (ER) stress (reflecting high secretion in mesenchymal cells) (*Appendix S5*, **Figure 3E**).

However, the cells exhibit considerable heterogeneity, seen most clearly by comparing the cells in clusters 4 and 6, which vary in their expression signatures and in their fates (**Figure 4A,B and Figure S7**). While cells in both clusters are highly proliferative, cells in cluster 4 have begun to lose MEF identity, show lower ER stress, and have higher OKSM-cassette expression, while cells in cluster 6 have the opposite properties (**Figure 3D,E** and **Figure S6B**). The cells in the two clusters show clear differences in their enrichment in the ancestral distribution of iPSCs (**Figure 4D)**. The majority (54%) of the day 2 ancestors of iPSCs lie in cluster 4, while only a small fraction (3%) lie in cluster 6. Clusters 4 and 6 also show clear differences in their descendants (**Figure 4A,C** and **Figure S7A**): the descendants of cells in cluster 6 are strongly biased toward the Valley of Stress (*e.g.*, 81% of Cluster 6 cell descendants are in clusters 8-11 by day 8 *vs.* 18% for cluster 4), while cluster 4 is strongly biased toward the Horn of Transformation (*e.g.*, 81% in clusters 19-21 *vs*. 12% for cluster 6).

The strongest difference in gene expression between clusters 4 and 6 was seen for *Shisa8* (detected in 67% vs. 3% of cells in clusters 4 and 6, respectively) (**Figure 3F**, **Figure S6B**) and *Shisa8*^+^ cells are enriched among the day 2 ancestors of iPSCs (**Figure S6B**). Notably, *Shisa8* is strongly associated with the entire trajectory toward successful reprogramming (**Figure 3F**): it is expressed in the Horn, pre-iPSCs, and iPSCs, but not in the Valley or in the alternative fates of neurogenesis and placental development. The expression pattern of *Shisa8* is similar to, but stronger than, that of *Fut9* (**Figure S5**), a known early marker of successful reprogramming that synthesizes the surface glyco-antigen SSEA-1 [*12*]. *Shisa8* is a little-studied mammalian-specific member of the Shisa gene family in vertebrates, which encodes single-transmembrane proteins that play roles in development and are thought to serve as adaptor proteins [*48*]. Our analysis suggests that *Shisa8* may serve as a useful early predictive marker of eventual reprogramming success and may play a functional role in the process.

### Cells in the valley of stress induce a Senescence Associated Secretion Phenotype (SASP)

By day 4, cells display a bimodal distribution of properties that is strongly correlated with their eventual descendants: cells in cluster 8 (low proliferation, high MEF identity, **Figure 3D,E** and **Figure S6C**) have 95% of their descendants in the Valley (**Figure 4A,B** and **Figure S7A**), while cells in cluster 18 (high proliferation, low MEF identity, **Figure 3D,E** and **Figure S6C**) have 94% of their descendants in the Horn (**Figure 4A,B** and **Figure S7A** and **table S5**). Cells in cluster 7 show intermediate properties and have roughly equal probabilities of each fate (**Figure 4A,B** and **Figure S7A**).

Along the trajectory from cluster 8 to the Valley (days 10-16; **Figure 4A,B** and **4E,F**), cells show a strong decrease in cell proliferation (**Figure 3E**), accompanied by increased expression of various cell-cycle inhibitors, such as *Cdkn2a*, which encodes *p16*, an inhibitor of the *Cdk4/6* kinase and halts G1/S transition (**Figure 3F**), *Cdkn1a* (*p21*), and *Cdkn2b* (*p15*) (**Figure S6D**), which peaks in the Valley. The cells show increased expression of D-type cyclin gene *Ccnd2* **(Figure S5,S6D**) associated with growth arrest [*49*]. A subset of the cells in the Valley (29%; clusters 12 and 14) showed high activity for a gene module that is correlated with a *p53* proapoptotic signature, compared to all other cells inside the Valley (p-value< 10^−16^, average difference 0.17, *t*-test) and outside the Valley (p-value< 10^−16^, average difference 0.32, *t*-test) (**Figure 3E, Figure S6E**).

Cells in the Valley also show activation of signatures of extracellular-matrix (ECM) rearrangement and secretory functions (**Figure 3E, Figure S6E**). Because these properties are consistent with a senescence associated secretory phenotype (SASP), we used a SASP signature involving 60 genes [*50*] (*Appendix S5*). Cells with this signature appear on day 10 and continue through day 16, consistent with previous reports concerning the timing of onset of stress-induced senescence [*50*] (**Figure 3E, Figure S6E**).

SASP, which has key roles in wound healing and development that are relevant for reprogramming biology, includes the expression of various soluble factors (including *Il6*), chemokines (including *Il8*), inflammatory factors (including *Ifng*), and growth factors (including *Vegf*) that can promote proliferation and inhibit differentiation of epithelial cells [*50*]. Recent reports have suggested that secretion of *Il6* and other soluble factors by senescent cells can enhance reprogramming [*51*]. Although detectable levels of *Il6* mRNA were present in only a small fraction of cells both in 2i and serum (0.2%) at days 12 and 16 (0.34% in all cells), the overall SASP signature was evident in 72% of cells in the Valley (*vs*. 11% elsewhere, primarily in day 0 MEFs). This suggests that the senescent cells in the Valley are thus likely to have paracrine effects on cells that successfully emerge from the Horn. We return to this point below.

### Other cells at day 4 are strongly biased toward the Horn of Transformation

For the remaining cells at day 4, the forward trajectory is characterized by high proliferation and loss of MEF identity (**Figure 3B,E**), and the descendants are strongly biased toward the Horn at day 8 (**Figure 4A,B** and **Figure S7A** and **table S5**). The Horn is distinguished as a point of transformation, where cells that have lost their mesenchymal identity are beginning their transitions to an epithelial fate. As discussed below, a minority of cells in the Horn have begun to express activators of a pluripotency expression program.

Following Dox withdrawal and media replacement on day 8, the cells in the Horn adopt one of four alternative outcomes by day 12 (senescence, neuronal program, placental program, and pre-iPSCs). Roughly half appear to become senescent, migrating through clusters 19 and 10 to the Valley (**Figure 4A**). The fate of the remaining cells is strongly influenced by the culture medium. In serum conditions, the proportion of these cells that transition to neuronal, placental and pre-iPSC states is 62%, 13% and 26%, respectively. By contrast, the proportions in 2i condition are 3%, 37% and 59% (**table S5**). These results are consistent with the presence in the 2i medium of two small-molecule inhibitors to inhibit differentiation, including one reported to inhibit neuronal differentiation [*52*].

### Neuronal-like and placental-like cells arise during reprogramming

We next studied two unusual cell populations: placental-like cells (clusters 24 and 25, **Figure 3B,D** and **Figure 4A,B,E,F**) at day 12 and neural-like cells (clusters 26 and 27, **Figure 3B,D** and **Figure 4A,B,E,F**) at day 16. The first group was characterized by high activity of two gene modules enriched in signatures for “epithelial cell differentiation,” “placenta development,” and “reproductive structure development,” while the second group showed high activity of signature for “neuron differentiation,” “axon development,” and “regulation of nervous system development” (**table S2**, and **Figure 3B**,**C,E**). While an earlier study from our group [*45*] had noted that upregulation of lineage-specific genes from unrelated lineages can be detected during reprogramming, distinct cell populations representing these differentiated states have not previously been recognized and characterized, to the best of our knowledge.

Both populations showed a substantial decrease in proliferation (**Figure 3E, Figure S6E**). To explore if a common mechanism was responsible for this change, we examined 98 cell-cycle-related genes [*53*] to identify those that were differentially upregulated in the placenta and neural clusters compared to all other clusters. The most distinctive characteristic was the high expression of *Cdkn1c*, which encodes a cell-cycle inhibitor (*p57*) that promotes G1 arrest (**Figure 3F**) and is required for maintenance of some adult stem cells [*54*]. Other features are also shared between these two alternative lineages and adult stem cells–including the expression of *Lgr5*, a marker of adult epithelial stem cells in certain tissues [*55*] (**Figure S5**).

The neural-like cells reside in a large “spike” observed at day 16 in serum but not 2i conditions (16% vs. 0.1% of cells), presumably due to differentiation inhibitors in the latter conditions. Cells near the base of the spike (cluster 26, **Figure 3D** and **Figure 4E,F**) expressed neural stem-cell markers (including *Pax6* and *Sox2,* **Figure 3E**, **Figure S5**), while cells further out along the spike (cluster 27, **Figure 3D**) expressed markers of neuronal differentiation (including *Neurog2* and *Map2,* **Figure S5**). The cells thus appear to span multiple stages of neurogenesis along the length of the spike (**Figure 3E**).

Analysis of the developmental landscape suggests a potential mechanism for triggering neural differentiation. The ancestors of neural-like cells are largely found in cluster 23 on day 12 (**Figure 4A,F** and **Figure S7C** and **table S5**). At least 19% of cells in cluster 23 express *Cntfr*, an Il6-family receptor that plays a critical role in neuronal differentiation and survival [*56*] (**Figure 3F**); the true proportion is likely to be higher because the gene has low expression. Contemporaneously, senescent cells in the Valley at day 12 express activating ligands (*Crlf1* and *Clcf1*) of *Cntfr* (**Figure S5**). Thus, neural differentiation may be triggered by paracrine signals from senescent cells to Cntfr-expressing cells.

The placental-like cells express high levels of certain imprinted genes on chromosome 7 (*Cdkn1c, Igf2, Peg3, H19* and *Ascl2;* **Figure 3F**, **Figure S5**), as well as TFs (*Cdx2* and *Sox17*) associated with placental development [*57, 58*] (**Figure S5**). They also show elevated levels of an ER stress signature (**Figure 3E**), consistent with the secretory nature of placental cells and observations of placental cells *in vivo* [*59*]. We checked whether the placental-like cells resembled recently described extraembryonic endodermal (XEN) cells from an iPSC reprogramming study [*44*], but found that they do not share the distinctive XEN signature of the cells reported in that paper (*Appendix S5*). The proportion of cells in the placental-like population fell substantially from day 12 to day 16 in 2i condition, although our optimal-transport analysis could not confidently infer whether the decrease is due to cells dying, being overtaken by faster-growing cells, or transitioning to other fates (**Figure S4A**).

### Trajectory to successful reprogramming reveals a continuous program of gene activation

We next studied the trajectory leading to reprogramming (**Figure 4D,E**), which passes through pre-iPSCs (cluster 28; **Figure 4A,B**) at day 12 en route to iPSC-like cells at day 16. The iPSC-like cells in serum conditions (which reside in cluster 31) closely resemble fully reprogrammed cells grown in serum (cluster 32). By contrast, the iPSC-like cells under 2i conditions are spread across three clusters (cluster 29-31).While the cells in cluster 31 resemble fully reprogrammed cells grown in 2i (cluster 33), those in cluster 29 show distinct properties suggestive of partial differentiation. In particular, cluster 29 shows lower proliferation, lower *Nanog* expression, and increased expression of genes related to differentiation (**Figure 3D-F**).

In contrast to initial descriptions of reprogramming as involving two “waves” of gene expression, the trajectory of successful reprogramming reveals a more complex regulatory program of gene activity (**Figure 5A**). By grouping genes according to their temporal patterns of activation in cells on the OT-defined trajectory to successful reprogramming, we can obtain a rich collection of markers for particular stages (**Figure 5A**). In particular, we identified 47 genes that appear late in successfully reprogrammed cells (for example, *Obox6*, *Spic, Dppa4*); these may provide useful markers to enrich fully reprogrammed iPSCs (**table S6**).

**Figure 5.**
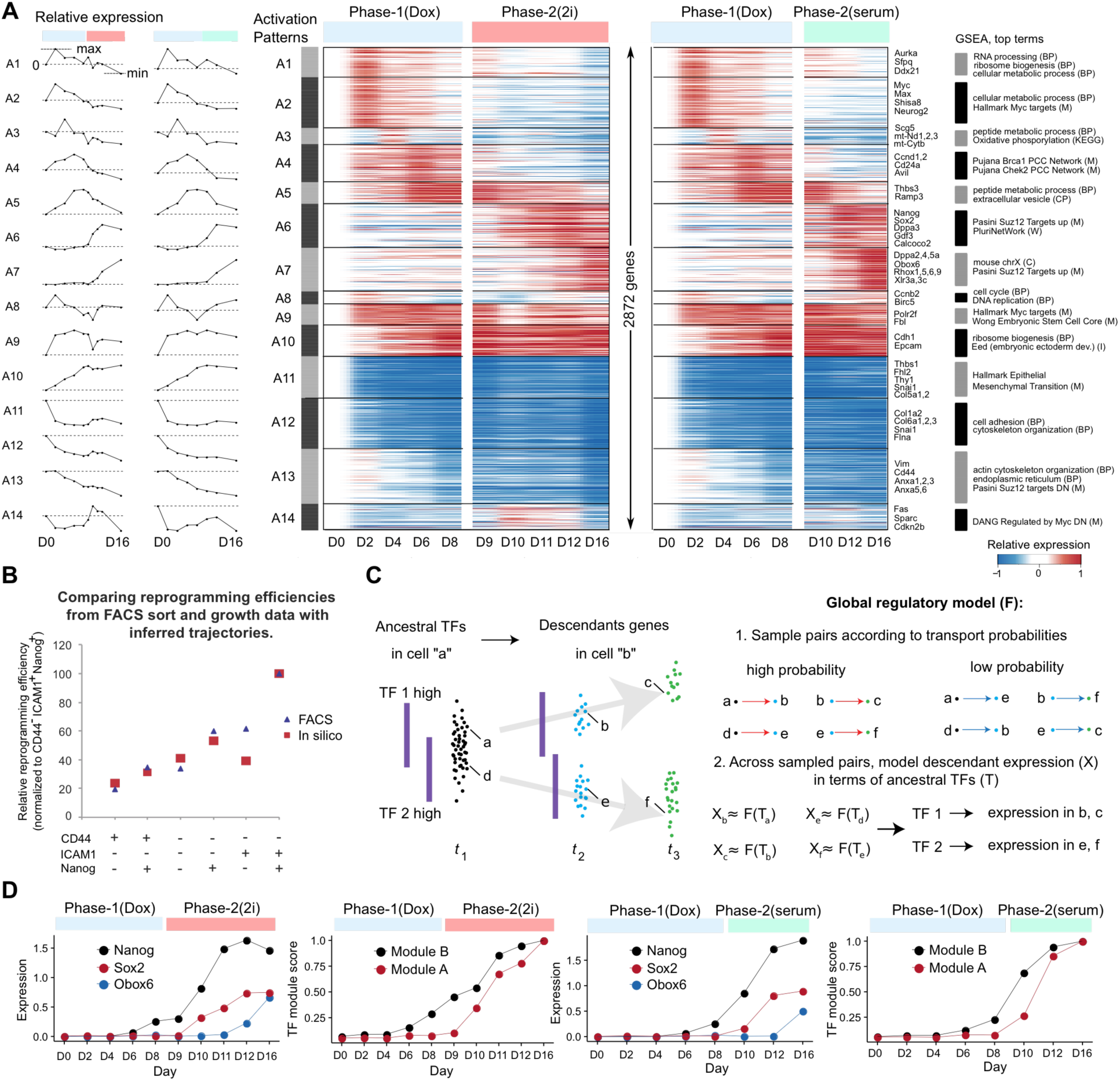
Gene activation along trajectory to successful reprogramming. (**A**) Classification of genes into 14 groups based on similar temporal expression profiles along the trajectory to successful reprogramming. Averaged gene expression profiles for each group, in 2i and serum conditions (left).Heatmap for genes within each group, with intensity of color indicating log2-fold change in expression relative to day 0 (middle). Representative genes and top terms from gene-set enrichment analysis for each group (right). **(B)** Comparison of FACS and *in* silico sorting experiments. Scatterplot shows reprogramming efficiencies determined by FACS sort and growth experiments (blue triangles) (*16*) and our computationally inferred trajectories (red squares). The specific cell surface markers used for the *in silico* and experimental methods are indicated. Reprogramming efficiencies for these categories (calculated both experimentally and *in silico*) are normalized to the percentage of EGFP+ colonies in CD44^-^ ICAM1^+^ Nanog^+^ condition (details found in *Appendix 5*). (**C**) Schematic of regulatory model in which TF expression in ancestral cells is predictive of gene expression in descendant cells. (**D**) Onset of iPSC-associated TFs in 2i (left) and serum (right). (Top) Mean expression levels weighted by iPSC ancestral distribution probabilities (Y axis) of *Nanog*, *Obox6*, and *Sox2* at each day (X axis). (Bottom) Normalized expression of TF modules “A” and “B” from our regulatory model (as in **B**) that were associated with gene expression in iPSCs. (see Figure S8 for details).

### Paracrine signaling from the Valley may influence late stages of reprogramming

The simultaneous presence of multiple cell types raises the possibility of paracrine signaling, with secreted factors from one cell type binding to receptors on another cell type. We noted one such potential interaction above, with SASP+ cells in the Valley secreting *Crlf1, Clcf1* and neural-llke cells on days 12 and 16 expressing the cognate receptor *Cntfr*.

To systematically identify potential opportunities for paracrine signaling, we defined an interaction score, I_A,B,X,Y,t_, as the product of (1) the fraction of cells in cluster A expressing ligand X and (2) the fraction of cells in cluster B expressing the cognate receptor Y, at time t. Using a curated list of 149 expressed ligands and their associated receptors (*Appendix S5*), we studied potential interactions between all pairs of clusters for each ligand-receptor pair, as well as the aggregate signal across all pairs and across those pairs related to the SASP signature.

The potential for paracrine signaling varied sharply across the time course, as well as across cell types. Potential interactions are initially high, as cells with MEF identity retain their secretory functions; drop dramatically by day 6 (**Figure S8A**), after cells have lost their MEF identity (**Figure 3B,C,E**); rise steadily from day 8 to day 11, as secretory cells in the Valley emerge; and then drop again from days 12 to 16, as the abundance of cells in the Valley decreases (**Figure S8A**). The same pattern is seen when considering only the 20 ligands in the SASP signature (**Figure S8B**).

Notably, potential interactions are observed between cells in the Valley and each of iPSC, neural-like and placental-like cells. At day 16, cells in the Valley (clusters 15 and 16) express SASP ligands, while iPSCs (clusters 29-33) express receptors for these ligands (**Figure S8C**), with the highest frequency seen for the chemokine *Cxcl12* and receptor *Dpp4* (**Figure S8D**). As noted above, at days 12 and 16, the ligands *Crlf1* and *Clcf1* cells are expressed in the Valley while their receptor *Cntfr* is expressed in the neural spike (**Figure 3E, Figure S8E**). The interaction between *Cntfr* and *Crlf1* is ranked as the top interaction among all ligand-receptor pairs (**Figure S8E**). Interestingly, at day 12, many placental-like cells express the ligand *Igf2* while cells in the Valley express receptors *Igf1r and Igf2r* (**Figure S8F**). We emphasize that the analysis can only nominate *candidate* interactions; experimental studies will be needed to determine if functional interactions actually occur.

### X-chromosome reactivation follows activation of early and late pluripotency genes

The reversal of X-chromosome inactivation in female cells is known to occur in the late stages of reprogramming and is an example of chromosome-wide chromatin remodeling. A recent study [*60*] reported that X-reactivation follows the activation of various pluripotency genes, based on immunofluorescence and RNA FISH in single cells. To assess X-reactivation, from scRNA-Seq data, we characterized each cell with respect to signatures of X-inactivation (*Xist* expression), X-reactivation (proportion of transcripts derived from X-linked genes, normalized to cells at day 0), and early and late pluripotency genes (*Appendix S5*). Along the trajectory to successful reprogramming (but not elsewhere, **Figure 3E**), cells at day 12 show strong downregulation of *Xist* but do not yet display X-reactivation. X-reactivation is complete at day 16, with the signature having risen from 1.0 to ∼1.6, consistent with the expected increase in X-chromosome expression [*61*]. Analysis of the trajectory confirms that activation of both early and late pluripotency genes precedes *Xist* downregulation and X-reactivation (*Appendix S5*).

### Some cell populations are enriched for aberrant genomic events

We next searched for other coherent increases or decreases in gene expression across large genomic regions, which might indicate the presence of copy-number variations (CNVs) in specific cells. Our approach built on methods we had previously developed for single-cell tumor analysis [*53*] (*Appendix S5*). We first looked for whole chromosome aberrations, and found that 0.9% of cells showed significant up- or down-regulation across an entire chromosome; the expression-level changes were largely consistent with gain or loss of a single chromosome (Appendix S5, **Figure A11A**). Next, we looked for evidence of large subchromosomal events, by analyzing regions spanning 25 consecutive housekeeping genes (median size ∼25 Mb). We found significant events in ∼0.8% of cells. The frequency was highest (2.8%) in cluster 14, consisting of cells in the Valley of Stress enriched for a DNA damage-induced apoptosis signature. The frequency was 2-to-3fold lower in other cells in the Valley (enriched for senescence but not apoptosis), in cells en route to the Valley (clusters 8 and 11), and in fibroblast-like cells at days 0 and 2. Notably, it was much lower (6-fold) in cells on the trajectory to successful reprogramming (*Appendix S5*, **Figure A11B,C**). Direct experimental evidence would be needed to confirm these events, and to clarify if the aberrations were preexisting in the MEF population, or if they accumulated during the course of reprogramming.

### Inferred trajectories agree with experimental results from cell sorting

To test the accuracy of the probabilistic trajectories calculated for each cell based on optimal transport, we compared results based on the trajectories to experimental data from a recent study of reprogramming of secondary MEFs [*16*]. In that study, cells were flow-sorted at day 10, based on the cell-surface markers CD44 and ICAM1 and a Nanog-EGFP reporter gene, and each sorted population was grown for several days thereafter to monitor reprogramming success. Gene expression profiles were obtained from each population at day 10 and CD44^-^ICAM1^+^Nanog^+^ population at day 15, together with mature iPSCs and ESCs. Reprogramming efficiency was lowest for CD44^+^ICAM^-^ Nanog^-^ cells, intermediate for CD44^-^ICAM1^+^Nanog^-^ and CD44^-^ICAM1^-^Nanog^+^ cells, and highest for CD44^-^ICAM1^+^Nanog^+^ cells.

We emulated the flow-sorting-and-growth protocol *in silico*, by partitioning cells based on transcript levels of the same three genes at day 10 and predicting the fates of each population at day 16 based on the inferred trajectory of each cell in the optimal transport model. Our computational predictions showed good agreement with these earlier experimental results (**Figure 5B**), with respect to both reprogramming efficiency and changes in gene-expression profiles. In particular, our *in silico* results showed 93% correlation with results from the earlier study concerning relative reprogramming efficiencies for six categories of sorted cells (*p* value= 0.0023) (**Figure 5B**, *Appendix S5*). Notably, the computationally inferred trajectory of doublepositive cells rapidly transitioned toward iPSCs and continued in this direction through the end of the time course (**Figure 5B**). Only one category (CD44-ICAM+Nanog-) differed significantly. Differences may reflect the fact that experimental protocols were not identical (*e.g.*, the earlier study [*16*] maintains continuous expression of OSKM and supplements the medium with an ALK-inhibitor and vitamin C).

### Inferring transcriptional regulators that control the reprogramming landscape

Our optimal transport map provides an opportunity to infer regulatory models, based on association between TF expression in ancestors and gene expression patterns in descendants. Specifically, we identified TFs by two approaches (**Figure 5C**): (i) a global regulatory model, to identify modules of TFs and target genes and (ii) enrichment analysis, to identify TFs in cells having many *vs*. few descendants in a target cell population of interest (*Appendix S5)*.

We first studied gene regulation along the trajectories to placental-like and neural-like cells (**Figure S9**). For placental-like cells, the analysis pointed to 22 TFs (**Figure S9A,B** and **table S7**). Of the four most enriched (*Pparg*, *Cebpa, Gcm1*, and *Gata2*), all have been reported to play roles in placenta development [*62*]. For example, *Gcm1* was detected in 42% of cells at day 10 with a high proportion (>80%) of descendants in the placental-like fate but only 0.7% of those cells with a low proportion (<20%) (57-fold enrichment). For neural-like cells, the analysis pointed to 10 TFs (*Pax3, Msx1, Msx3, Sox3, Sox11, Tal2, En1, Foxa2, Gbx2,* and *Foxb1*). All have been implicated in various aspects of neural development (**Figure S9C**, *Appendix S5*) [*62-70*].

Next, we focused on identifying TFs that play roles along the trajectory to successful reprogramming (**Figure 5D** and **Figure S9D,E**). The global regulatory model generated two regulatory modules, A and B, with 61 TFs in module A, 16 in module B, and 11 in both (**Figure S9D,E**). Module A involves target genes active across clusters 29-31, while Module B involves target genes that are more active in cluster 31, which contains more fully reprogrammed cells. The TFs in these modules are progressively activated across the trajectory of successful reprogramming. For Module B, the TFs are active in 13% of cells in the Horn on day 8, while target-gene activity is evident (at >80% of the levels observed in iPSCs) in 1.3%, 10%, and 21% of their descendant cells in days 10, 11, and 12 in 2i conditions; the pattern in serum conditions is similar, although with lower overall frequency (11% of cells by day 12). The onset of TFs and target genes in Module A lags by 1-2 days (**Figure 5D**).

To identify TFs likely to play a key role in the final stages of reprogramming, we used enrichment analysis to identify TFs enriched in cells at day 12 with a high vs. low proportion (>80% vs. <20%) of successfully reprogrammed descendants and then focused on the intersection of this set with the 66 TFs from the global regulatory analysis above.

The analysis pointed to 9 TFs associated with a high probability of success in the late stages of reprogramming (**Figure S9F**). Of these, five (*Sox2, Nanog, Hesx1, Esrrb, Zfp42*) have established roles in regulation of pluripotency [*71-73*], while the remaining four (*Obox6, Spic, Mybl2,* and *Msc*) have not previously been implicated. Among these novel factors, *Obox6* stands out as having the greatest enrichment in highvs. low-probability cells (68-fold, 9.3% vs ∼0.14%) **(Figure S9F)**.

### Forced expression of *Obox6* enhances reprogramming

Finally, we experimentally studied *Obox6*, which the regulatory analysis discovered to be strongly correlated with reprogramming success. *Obox6* (oocyte-specific homeobox 6) is a homeobox gene of unknown function that is preferentially expressed in the oocyte, zygote, early embryos and embryonic stem cells [*74*].

To test whether *Obox6* also plays an active role in the process, we investigated whether expressing *Obox6* along with OKSM during days 0-8 can boost reprogramming efficiency. We infected our secondary MEFs with a Dox-inducible lentivirus carrying either *Obox6*, the known pluripotency factor *Zfp42* [*73*], or no insert as a negative control. Both *Obox6* and *Zpf42* increased reprogramming efficiency of secondary MEFs by ∼2-fold in 2i and even more so in serum, with the result confirmed in multiple independent experiments (**Figure 6A** and **B,** and **Figure S10**). Assays in primary MEFs showed similar increases in reprogramming efficiency (**Figure S10**). These results highlight *Obox6* as meriting further study in the context of reprogramming.

**Figure 6.**
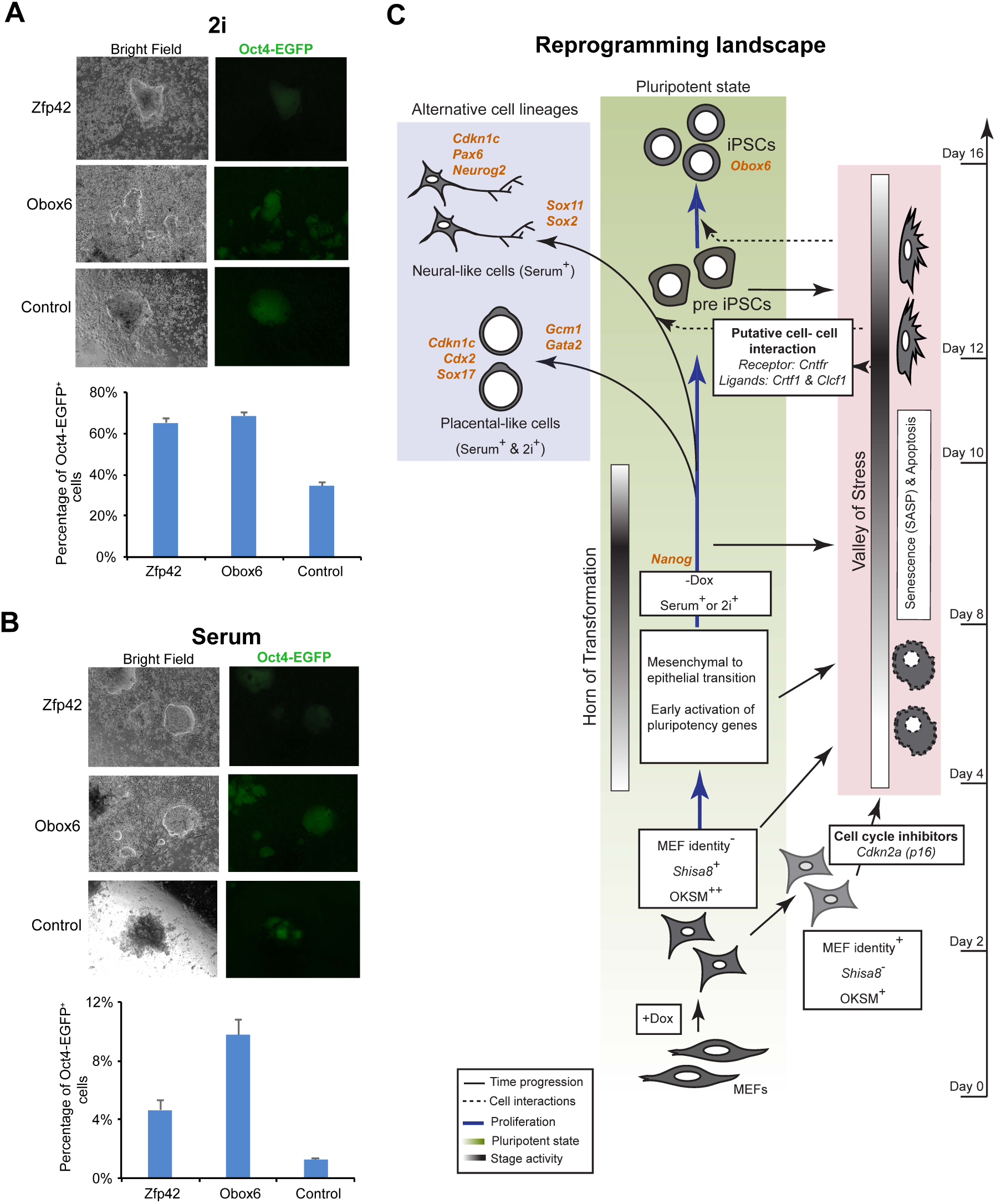
Effect of overexpression of *Obox6* and *Zpf42* on reprogramming efficiency in secondary MEFs. **(A)** Bright field and fluorescence images of iPSC colonies generated by lentiviral overexpression of *Oct4*, *Klf4*, *Sox2*, and *Myc* (OKSM) with either an empty control, *Zfp42* or *Obox6* expression cassette, in either Phase-1(Dox)/Phase-2(2i) and Phase-1(Dox)/Phase-2(serum) conditions (indicated). Cells were imaged at day 16 to measure Oct4-EGFP^+^ cells. Bar plots representing average percentage of Oct4-EGFP^+^ colonies in each condition on day 16 are included below the images. Shown are data from one of five independent experiments, with three biological replicates each. Error bars represent standard deviation for the three biological replicates. **(B)** Schematic of the overall reprogramming landscape highlighting: the progression of the successful reprogramming trajectory, alternative cell lineages, and specific transition states (Horn of Transformation). Also highlighted are transcription factors (orange) predicted to play a role in the induction and maintenance of indicated cellular states, and putative cell-cell interactions between contemporaneous cells in the reprogramming system.

## Discussion

Large-scale profiling of single cells has opened new prospects for systematically dissecting the processes and programs underlying cellular differentiation. While the ideal approach would involve profiling the *same* cells across many time points, current methods for collecting rich information (such as gene expression profiles) are destructive. An alternative approach would be to mark cells with unique barcodes at a time *t* and sample cells at subsequent time points to capture multiple descendants with the same barcode at multiple subsequent time points.However, the approach can only detect fates that were already determined at time *t* and is not well suited for studying events, as here, that occur over short time periods and few cell divisions, due to the small number of descendant cells with each barcode available for sampling.

Here, we describe an approach for studying developmental trajectories that requires sampling a time-dependent distribution across a time series and using optimal transport with growth to couple the distributions. For any set S of cells, one can use this approach to (i) infer a distribution over potential ancestors and descendants; (ii) study trajectories to and from S; and construct regulatory models for the trajectory. The approach aims to capture the flexibility inherent in Waddington’s metaphor of marbles rolling in a developmental landscape (vs. his metaphor of railroad tracks) and thus to loosen current constraints on studying developmental potential.

Based on this framework we developed WADDINGTON-OT for exploratory data analysis of developmental landscapes. WADDINGTON-OT enables analyses (optimal-transport analysis, gene clustering, cell clustering, regulatory analysis) and visualization of features (such as time, gene-expression levels, gene-signature levels, and trajectories). With these tools, one can easily measure the extent to which any single gene — or, more powerfully, gene signatures — serves not only as a reliable marker of a cellular state but also as a predictor of a subsequent cellular fate. Gene signatures can be used to sort classes of cells *in silico* and analyze the events along their trajectories, or to guide the design of laboratory reagents, such as reporter constructs and antibodies, to do the same in the laboratory.

To test these ideas, we applied WADDINGTON-OT to study cellular reprogramming of fibroblasts to iPSCs. The analysis (**Figure 6C**) recapitulates known features of reprogramming, as well as providing new insights including:

(1) *discovering heterogeneity within known cell populations*. For example, cells at day 2 already show clear differences in proliferation rates, MEF identity and a gene signature predictive of reprogramming success.

(2) *identifying new cell populations.* For example, the analysis reveals distinct neural- and placental-like cell populations that represent reprogramming failures.

(3) *disentangling cellular trajectories*. For example, data from a heterogeneous population can be decomposed into simultaneous trajectories toward distinct destinations, such as senescence, neural differentiation, and iPSCs.

(4) *discovering genes and gene signatures associated with specific states and predictive of specific fates.* For example, *Shisa8* and *Obox6* are identified as early and late markers of reprogramming success, respectively.

(5) *associating regulators with specific trajectories.* For example, TFs likely to play key regulatory roles are identified along the neural, placental, and iPSC pathways, either predicting later changes in gene expression or maintaining stable states.

(6) *prediction of interactions between different contemporaneous cells*. For example, the data highlight potential effects of factors secreted by senescent cells on more proliferative cells.

Several lines of evidence support the validity of the approach: (i) the analysis directly discovers many known aspects of reprogramming and, where new processes and cell types are identified, they are consistent with previous knowledge; (ii) TFs discovered by the regulatory analysis of neural-like, placental-like and iPSCs are consistent with prior knowledge; (iii) experimental tests support the identification of *Obox6* as a novel TF involved in reprogramming; and (iv) computational tests show that the biological results are robust to details of the analysis (e.g., choice of parameters, cost functions, growth rates and subsampling of the data).

Nonetheless, the methodology is clearly still at an early stage. Biological insights will likely emerge from applying the method to a range of biological systems. For example, normal development and physiology may show a greater role for cell-cell interactions than engineered reprogramming. Further insights will come by applying the method to additional types of singlecell characterization beyond gene expression, including epigenomic, proteomic, and imaging information.

There are a number of open mathematical issues, including how best to: optimize the choice and sampling depth of time points; define the distance function in gene-expression space; use information about cell proliferation rates; predict unobserved time points and choose the most informative ones to measure; incorporate lineage-tracing information where available; quantify uncertainty in transport maps; identify and characterize branching events leading to bimodal fate distributions; predict the effect of perturbing regulators; and smooth data over time (*Appendix S1*). Our framework currently assumes that cell fates depend only on internal variables, but we could readily incorporate cell-extrinsic influences in terms of bulk concentrations of secreted factors. Additionally, our framework exploits the assumption that trajectories are memoryless— that is, that a cell’s fates depends only on its current state rather than its history. Some processes are likely not memoryless—for example, due to aspects of a cell’s epigenomic state that are not fully captured in the gene expression profile. It will be important to develop methods to recognize such circumstances and incorporate them in the analysis.

Finally, improved experimental methods are needed to validate the many specific transition probabilities inferred by optimal transport. For now, the best way to test the inferences may be to compare it to results from denser temporal sampling, or to perturb predicted regulators and observe their effects on the maps.

In summary, the notion of developmental landscapes can provide not only a valuable metaphor but a precise analytical framework. The optimal-transport approach should provide insight into a wide range of biological problems, including organismal development, long-term physiological responses, and induced reprogramming.

## Acknowledgements

We thank M. Kowalczyk, D. Przybylski, S. Markoulaki, J. Drotar, D. Rooney, R. Flannery for technical advice and reagents; and J. Buenrostro for discussions. This work was supported by funds from Broad Institute (E.S.L, A.R.), NIH grants HD045022, R01 MH104610-15, and R01NS088538 (R.J.), and R01HD058013 (K.H.). J.S. is supported by the Helen Hay Whitney Foundation. G.S., M.T., and B.C. are supported by Klarman Cell Observatory at Broad Institute. J.S. and E.S.L. conceived and designed the study of reprogramming in single-cell resolution. G.S. and E.S.L. conceived the application of optimal transport; P.R. and A.R. provided input on the development of the approach. M.T., G.S., and B.C. developed WADDINGTON-OT. M.T., G.S., B.C., E.S.L. and A.R. analyzed the data, with assistance from J.C. All experiments were designed and performed by J.S., with input from R.J. and assistance from A.S., S.L, S.L., P.B., L.L., J.B., K.H. and V.S. The manuscript was written by E.S.L, A.R., G.S., B.C., M.T., and J.S.

## Supplementary Materials

### I. Appendix S1

#### Learning Developmental Processes with Optimal Transport

This appendix introduces a mathematical model for developmental processes, and a way to estimate them from data with optimal transport. This appendix is written for a mathematical audience. The appendix is organized as follows: section 1 reviews the concept of gene expression space and introduces our probabilistic framework for time series of expression profiles. Section 2 shows how we can compute an optimal coupling between adjacent time points by solving a convex optimization problem. Section 3 describes what we can do with transport maps. Specifically, section 3.1 shows how to compute origins and fates of cells, and section 6 shows how we learn gene regulatory networks to summarize the trajectories. Section 4 discusses a smoothed, continuous-time version of the method that interpolates between time points.

##### 1. Developmental processes in gene expression space

A collection of mRNA levels for a single cell is called an *expression profile* and is often represented mathematically by a vector in *gene expression space*. This is a vector space that has a dimension corresponding to each gene, with the value of the *i*th coordinate of an expression profile vector representing the number of copies of mRNA for the *i*th gene. Note that real cells only occupy an integer lattice in gene expression space (because the number of copies of mRNA is an integer), but we pretend that cells can move continuously through a real-valued *G* dimensional vector space.

As an individual cell changes the genes it expresses over time, it moves in gene expression space and describes a trajectory. As a population of cells develops and grows, a *distribution* on gene expression space evolves over time. When a single cell from such a population is measured with single cell RNA sequencing, we obtain a noisy estimate of the number of molecules of mRNA for each gene. We represent the measured expression profile of this single cell as a sample from a probability distribution on gene expression space. This sampling captures both (a) the randomness in the single cell RNA sequencing measurement process (due to subsampling reads, technical issues, etc.) and (b) the random selection of a cell from a population. We treat this probability distribution as *nonparametric* in the sense that it is not specified by any finite list of parameters.

In the remainder of this section we introduce a precise mathematical notion for a *developmental process* as a generalization of a stochastic process. Our primary goal is to infer the ancestors and descendants of subpopulations evolving according to an unknown developmental process. We claim this is possible over short time scales because it is reasonable to assume that cells don’t change too much and therefore we can infer which cells go where. We show in the remainder of this appendix how to do this with *optimal transport.*

###### 1.1 A mathematical model of developmental processes

We begin by formally defining a precise notion of the developmental trajectory of an individual cell and its descendants. Intuitively, it is a continuous path in gene expression space that bifurcates with every cell division. Formally, we define it as follows:

###### Definition 1

(single cell developmental trajectory). *Consider a cell x*(0) ∈ ℝ^*G*^*. Let k*(*t*) ≥ 0 *specify the number of descendants at time t, where k*(0) = 1*. A single cell developmental trajectory is a continuous function*

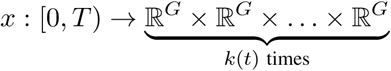

*This means that x*(*t*) *is a k*(*t*)*-tuple of cells, each represented by a vector in* ℝ^*G*^:

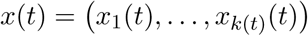

*We refer to the cells x*_1_(*t*)*, …, x*^*k*(*t*)^(*t*) *as the* ***descendants*** *of x*(0).

Note that we cannot directly measure the temporal dynamics of an individual cell because scRNASeq is a destructive measurement process: scRNA-Seq lyses cells so it is only possible to measure the expression profile of a cell at a single point in time. As a result, it is not possible to directly measure the descendants of that cell, and it is (usually) not possible to directly measure which cells share a common ancestor with ordinary scRNA-Seq. Therefore the full trajectory of a specific cell is unobservable.However, one can hope to learn something about the probable trajectories of individual cells by measuring snapshots from an evolving population.

Published methods typically represent the aggregate trajectory of a population of cells with a graph. While this recapitulates the branching path traveled by the descendants of an individual cell, we argue it over-simplifies the stochastic nature of developmental processes. Individual cells have the potential to travel through different paths, but in reality any given cell travels one and only one such path. Our goal is to describe this potential, which might not be a represented by a graph as a union of one dimensional paths.

Instead, we define a developmental process to be a time-varying *distribution* on gene expression space. We use the word distribution to refer to an object that assigns mass to regions of ℝ^*G*^. Note that we make a distinction between distribution and *probability* distribution, which necessarily has total mass 1. Distributions are formally defined as generalized functions (such as the delta function *δ*_*x*_) that act on test functions, but this is beyond our scope. For our purposes, a distribution is the same as a measure, but these notes don’t require a background in measure theory. One simple example of a distribution of cells is that we can represent a set of cells *x*_1_*, …, x*_*n*_ by the distribution

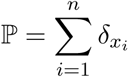

Similarly, we can represent a set of single cell trajectories *x*_1_(*t*)*, …, x*_*n*_(*t*) with a distribution over trajectories. This is a special case of a developmental process.

###### Definition 2

(developmental process). *A developmental process* ℙ_*t*_ *is a time-varying distribution on gene expression space.*

A developmental process generalizes the definition of *stochastic process*. A developmental process with total mass 1 for all time is a (continuous time) stochastic process, i.e. an ordered set of random variables with a particular dependence structure. Recall that a stochastic process is determined by its temporal dependence structure, i.e. the coupling between random variables at different time points. The coupling of a pair of random variables refers to the structure of their joint distribution.

The notion of coupling for developmental processes is the same as for stochastic processes, except with general distributions replacing probability distributions:

###### Definition 3

(coupling and transport map). *A coupling of a pair of distributions* ℙ, ℚ *on* ℝ^*G*^ *is a distribution π on* ℝ^*G*^ × ℝ^*G*^ *with the property that π has* ℙ *and* ℚ*as its two marginals. A coupling is also called a transport map.*

As a distribution on the product space ℝ^*G*^ *×*ℝ^*G*^, a transport map *π* assigns a number *π*(*A, B*) to any pair of sets *A, B ⊂*ℝ^*G*^

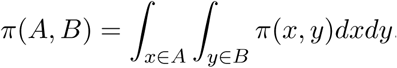

When *π* is the coupling of a developmental process, this number *π*(*A, B*) represents the mass transported from *A* to *B* by the developmental process. This is the amount of mass coming from *A* and going to *B*. When we don’t specify a particular destination, the quantity *π*(*A,*) specifies the full distribution of mass coming from *A*. We refer to this action as *pushing A* through the transport map *π*. More generally, we can also push a *distribution μ* forward through the transport map *π* via integration

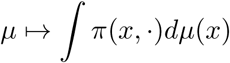

We refer to the reverse operation as pulling a set *B* back through *π*. The resulting distribution *π*(*, B*) encodes the mass ending up at *B*. We can also pull distributions *μ* back through *π* in a similar way:

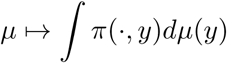

We sometimes refer to this as *back-propagating* the distribution *μ* (and to pushing *μ* forward as *forward propagation*).

Recall that a stochastic process is Markov if the future is independent of the past, given the present. Equivalently, it is fully specified by its couplings between pairs of time points. A general stochastic processes can be specified by further higher order couplings, but that is beyond our scope. We consider Markov developmental processes, which are defined in the same way:

###### Definition 4

(Markov developmental process). *A Markov developmental process* ℙ_*t*_ *is a time-varying distribution on* ℝ^*G*^ *that is completely specified by couplings between pairs of time points.*

It is an interesting question to what extent developmental processes are Markov. On gene expression space, they are likely not Markov because, for example, the history of gene expression can influence chromatin modifications, which may not themselves be reflected in the observed expression profile but could still influence the subsequent evolution of the process. However, it is possible that developmental processes could be considered Markov on some augmented space.

We now define the descendants and ancestors of subgroups of cells evolving according to a Markov developmental process. We extend our earlier definition of descendants as follows:

###### Definition 5

(descendants in a Markov developmental process). *Consider a set of cells S* ⊂ ℝ^*G*^*, which live at time t*_1_ *are part of a population of cells evolving according to a Markov developmental process* ℙ_*t*_*. Let π denote the transport map for* ℙ_*t*_ *from time t*_1_ *to time t*_2_*. The descendants of S at time t*_2_ *are obtained by pushing S through the transport map π.*

Note that if a developmental process is not Markov, then the descendants of *S* are not well defined. The descendants would depend on the cells that gave rise to *S*, which we refer to as the *ancestors* of *S*.

###### Definition 6

(ancestors in a Markov developmental process). *Consider a set of cells S* ⊂ ℝ^*G*^*, which live at time t*_2_ *and are part of a population of cells evolving according to a Markov developmental process* ℙ_*t*_*. Let π denote the transport map for* ℙ_*t*_ *from time t*_2_ *to time t*_1_*. The ancestors of S at time t*_1_ *are obtained by pushing S through the transport map π.*

Having defined the notion of a developmental process and the ancestors and descendants of subpopulations, we now turn to estimating these from data.

###### 1.2 Empirical developmental processes

Our main goal is to track the evolution of a developmental process from a scRNA-Seq time course. Suppose we are given input data consisting of a sequence of sets of single cell expression profiles, collected at *T* different time slices of development. Mathematically, this time series of expression profiles is a sequence of sets *S*_1_*, …, S*_*T*_ *⊂*ℝ^*G*^ collected at times *t*_1_*, …, t*_*T*_ *∈*ℝ.

###### Definition 7

(developmental time series). *A developmental time series is a sequence of samples from a developmental process* ℙ_*t*_ *on* ℝ^*G*^*. This is a sequence of sets S*_1_*, …, S*_*N*_ *⊂*ℝ ^*G*^*. Each S*_*i*_ *is a set of expression profiles in* ℝ ^*G*^ *drawn i.i.d from the probability distribution obtained by normalizing the distribution* ℙ_*t*_*i to have total mass* 1.

From this input data, we form an empirical version of the developmental process. Specifically, at each time point *t*_*i*_ we form the empirical probability distribution supported on the data *x ∈S*_*i*_. We summarize this in the following definition:

###### Definition 8

(Empirical developmental process). *An empirical developmental process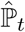 is a time varying distribution constructed from a developmental time course S*_1_*, …, S*_*N*_:

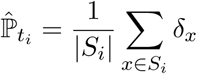

*The empirical developmental process is undefined for t ∉/ {t*_1_*, …, t*_*N*_ *}.*

Our goal is to recover information about a true, unknown developmental process ℙ_*t*_ from the empirical developmental process 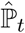. The measurement process of single cell RNA-Seq destroys the coupling, and the empirical developmental process we observe does not come with an informative coupling between successive time points. Over short time scales, we claim that it’s reasonable to assume that cells don’t change too much and therefore we can infer which cells go where and estimate the coupling.

In the next section, we propose to do this with *optimal transport*: we select the transport map *π* that minimizes the total work required for redistributing 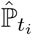 to 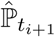 One motivation for minimizing this objective, is a deep relationship between optimal transport and dynamical systems that provides a direct connection to Waddington’s landscape: the optimal transport problem can formulated as a *least-action advection* of one distribution into another according to an unknown velocity field (see Theorem 1 in Section 6 below). At a high level, differentiation follows a velocity field on gene expression space, and the potential inducing this velocity field is in direct correspondence with Waddington’s landscape^1^. (See also **fig. A3**).

**Figure A1.**
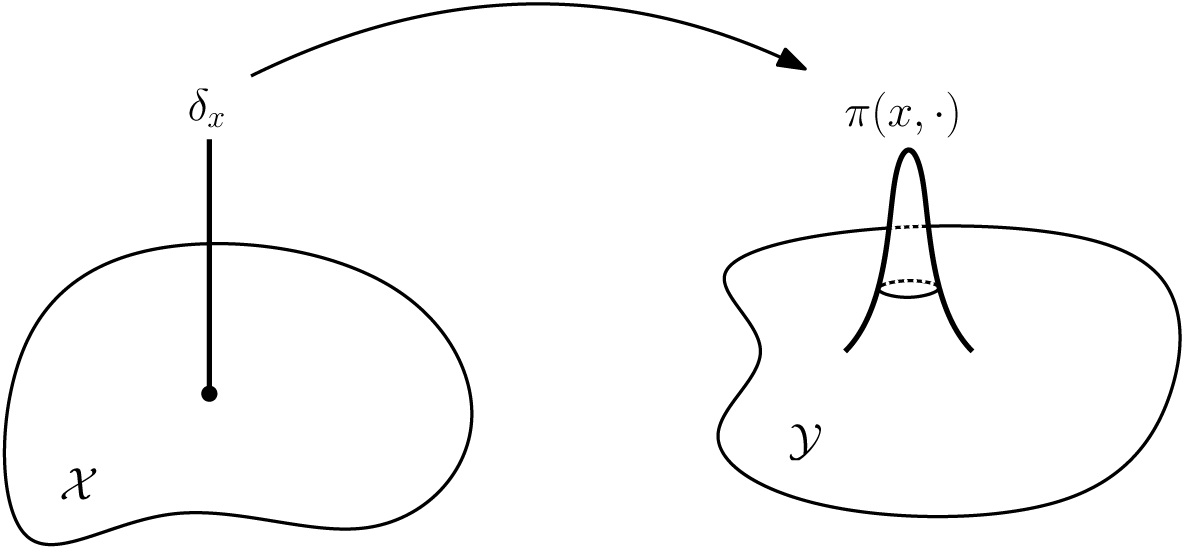
A transport plan *π* sends a unit mass *δ*_*x*_ to *π*(*x, ·*).

##### 2 Optimal transport for scRNA-Seq time series

In this section we describe how to compute probabilistic flows from a time series of single cell gene expression profiles by using optimal transport (S1). We show how to compute an optimal *coupling* of adjacent time points by solving a convex optimization problem.

Optimal transport defines a metric between probability distributions; it measures the total distance that mass must be transported to transform one distribution into another. For two measures ℙand ℚ on ℝ^*G*^, a *transport plan* is a measure on the product space ℝ^*G*^ × ℝ^*G*^ that has marginals ℙ and ℚ. In probability theory, this is also called a *coupling*. Intuitively, a transport plan *π* can be interpreted as follows: if one picks a point mass at position *x*, then *π*(*x,*) gives the distribution over points where *x* might end up (see **fig. A1**, and definition 3).

If *c*(*x, y*) denotes the cost^2^ of transporting a unit mass from *x* to *y*, then the expected cost under a transport plan *π* is given by

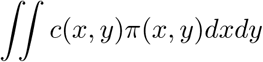

The *optimal* transport plan minimizes the expected cost subject to marginal constraints: minimize subject to

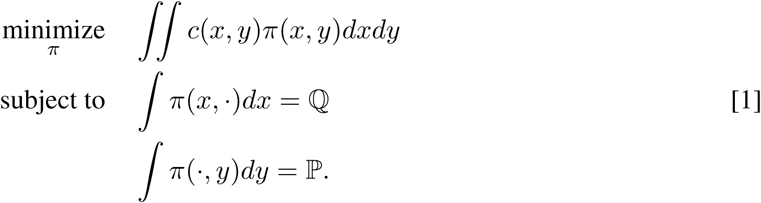

Note that this is a linear program in the variable *π* because the objective and constraints are both linear in *π*. Note that the optimal objective value defines the *transport distance* between ℙ and ℚ (it is also called the Earthmover’s distance or Wasserstein distance).Unlike most other ways to compare distributions (such as KL-divergence or total variation), optimal transport takes the geometry of the underlying space into account. For example, the KL-Divergence is infinite for any two distributions with disjoint support, but the transport distance between two unit masses depends on their separation.

When the measures ℙ and ℚ are supported on finite subsets of ℝ^*G*^, the transport plan is a matrix whose entries give transport probabilities and the linear program above is finite dimensional. In our setting, we form *empirical distributions* from the sets of samples *S*_1_*, …, S*_*T*_:

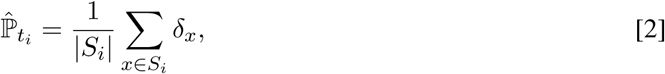

were *δ*_*x*_ denotes the Dirac delta function centered at *x* ∈ ℝ^*G*^. These empirical distributions 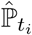 are finitely supported, and so we could solve the linear program [1] with 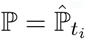 and 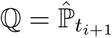.

However, the classical formulation [1] does not allow cells to grow (or die) during transportation(because it was designed to move piles of dirt and conserve mass). When the classical formulation is applied to a time series with two distinct subpopulations proliferating at different rates^3^, the transport map will artificially transport mass between the subpopulations to account for the relative proliferation. Therefore, we modify the classical formulation of optimal transport in equation [1] to allow cells to grow at different rates.

We assume that a cell’s measured expression profile *x* determines its growth rate *g*(*x*). This is reasonable because many genes are involved in cell proliferation (e.g. cell cycle genes). We assume *g*(*x*) is a known function (based on knowledge of gene expression) representing the exponential increase in mass per unit time, but also note that we ultimately allow the growth rate to be misspecified by leveraging techniques from *unbalanced transport* (S2). In practice, we define *g*(*x*) in terms of the expression levels of genes involved in cell proliferation. (See Appendix S2).

###### Derivation of transport with growth

For any cell *x*∈ *S*_*i-*1_, let *r*(*x, y*) be the fraction of *x* that transitions towards *y*. Then the amount of probability mass from *x* that ends up at *y* (after proliferation)is

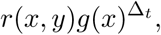

where Δ_*t*_ = *t*_*i*+1_ *- t*_*i*_. The total amount of mass that comes from *x* can be written two ways:

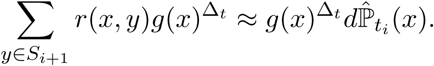

This gives us our first constraint. Similarly, we also have the constraint that *the total mass we observe at y is equal to the sum of masses coming from each x and ending up at y*. In symbols,

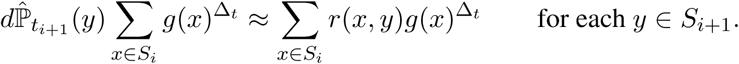

The factor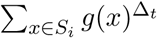on the left hand side accounts for the overall proliferation of all the cells from*S*_*i*_. Note that this factor is required so that the constraints are consistent: when one sums up both sides of the first constraint over *x*, this must equal the result of summing up both sides of the second constraint over *y*. Finally, for convenience, we rewrite these constraints in terms of the optimization variable

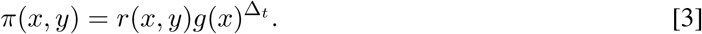

Therefore, to compute the transport map between the empirical distributions of expression profiles observed at time *t*_*i*_ and *t*_*i*+1_, we set up the following linear program:

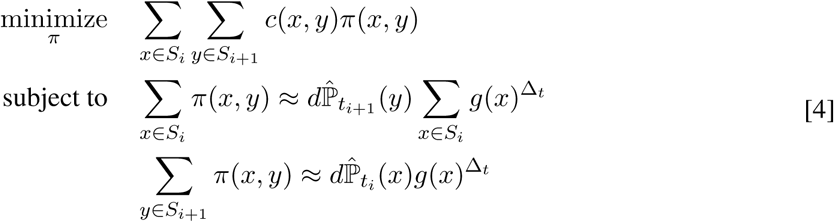

###### Regularization and algorithmic considerations

Fast algorithms have been recently developed to solve an entropically regularized version of the transport linear program (S3). Entropic regularization means adding the entropy 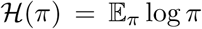 to the objective function, which penalizes deterministic transport plans (a purely deterministic transport plan would have only one nonzero entry in each row). Entropic regularization speeds up the computations because it makes the optimization problem strongly convex, and gradient ascent on the dual can be realized by successive diagonal matrix scalings (S3). These are very fast operations! This scaling algorithm has also been extended to work in the setting of *unbalanced transport*, where equality constraints are relaxed to bounds on KL-divergence (S2). This allows the growth rate function *g*(*x*) to be misspecified to some extent.

We use both entropic regularization and unbalanced transport. To compute the transport map between the empirical distributions of expression profiles observed at time *t*_*i*_ and *t*_*i*+1_, we solve the following optimization problem:

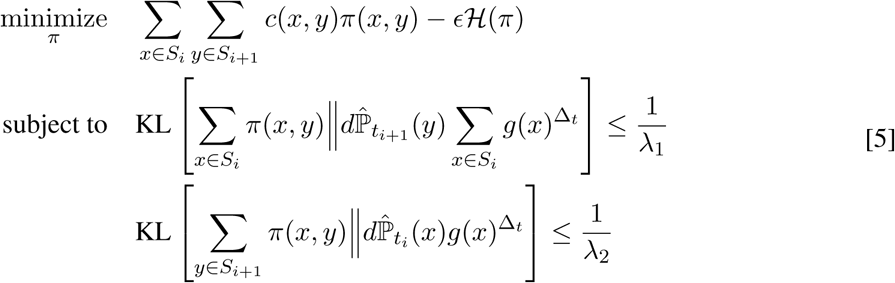

where ∈*, λ*_1_ and *λ*_2_ are regularization parameters (we describe our parameter selection strategy for the reprogramming dataset in Appendix 2). This is a convex optimization problem in the matrix variable *π* ∈ ℝ ^*N*^*i*^*×N*^*i*+1, where *N*_*i*_ = *S*_*i*_ is the number of cells sequenced at time *t*_*i*_. It takes about 5 seconds to solve this unbalanced transport problem using the scaling algorithm of Chizat et al. 2016 (S2) on a standard laptop with *N*_*i*_ ≈ 5000.

Note that the densities (on the discrete set *S*_*i*_) of the empirical distributions specified in equation [2] are simply 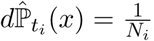. However, in principle one could use nonuniform empirical distributions (e.g.if one wanted to include information about cell quality).

To summarize: given a sequence of expression profiles *S*_1_*, …, S*_*T*_, we solve the optimization problem [5] for each successive pair of time points *S*_*i*_*, S*_*i*+1_. This gives us a sequence of transport maps as illustrated in **fig. A2**.

##### 3. What can we do with transport maps?

In the previous section we introduced the principle of optimal transport for time series of gene expression profiles. Given a time series of expression profiles *S*_1_*, …, S*_*T*_, we use this principle to compute a sequence of transport maps between subsequent time slices. In this section we show what we can learn from this sequence of transport maps. First, in section 3.1 we describe how to compute the *ancestors* and *descendants* of any subset of cells from this sequence of transport maps. Then, in section 6 we show how to fit gene regulatory networks that explain the flow of probability prescribed by the transport maps.

###### 3.1 Computing ancestors and descendants

For a specific set of cells, we refer to the contributions of cells from previous time points to as the *ancestors* of. Intuitively, we estimate the probable ancestors (or descendants) of by using the transport maps to propagate probabilities backward (or forward) in time.

**Figure A2.**
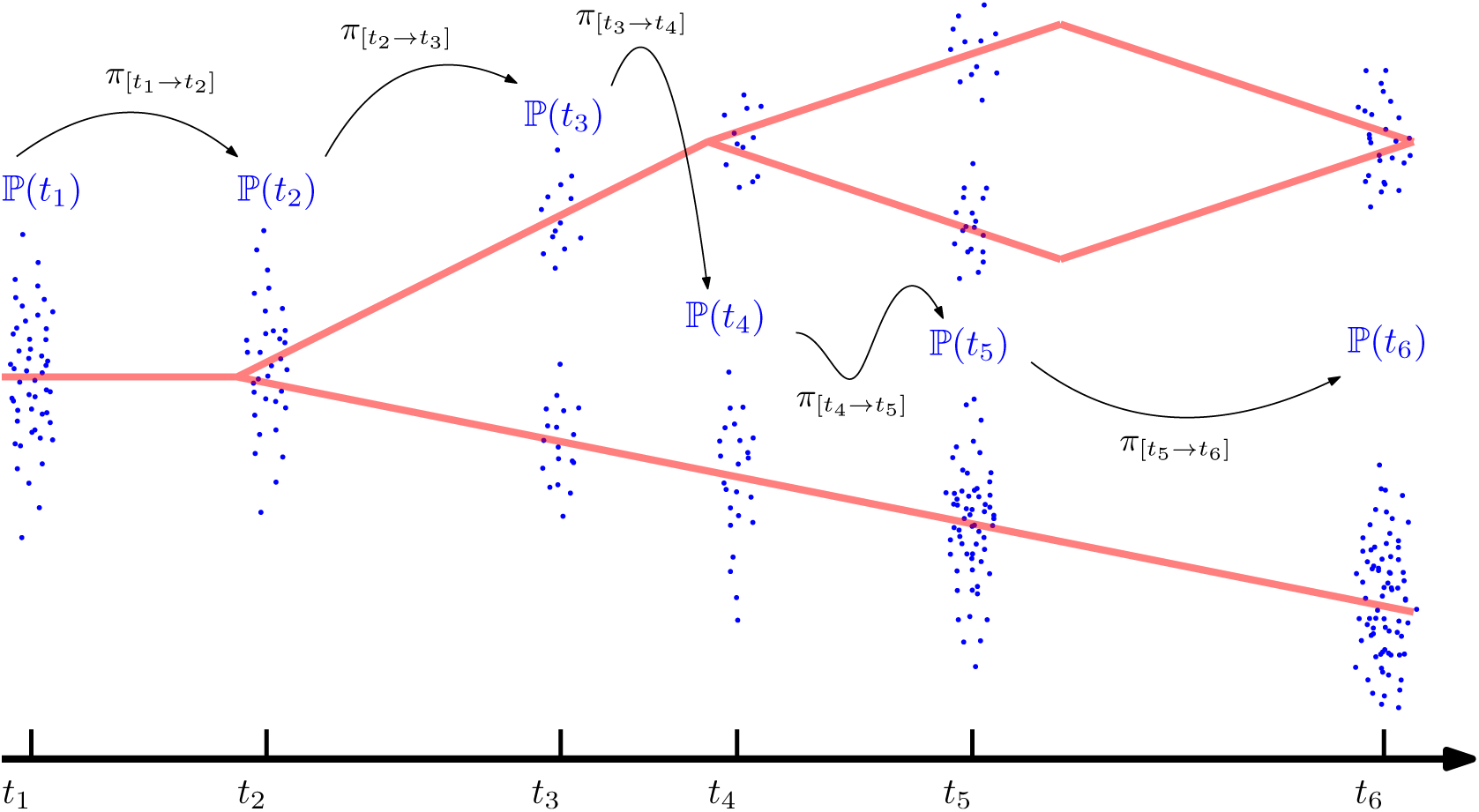
A diagram showing data *S*_*i*_ from a generic branching developmental process. The x-axis represents time and the y-axis represents expression. We compute transition probabilities *π* via optimal transport.

To make this more precise, we consider a single cell *y ∈ S*_*i*_. The column *π*(*·, y*) of the transport map *π* from *t*_*i-*1_ to *t*_*i*_ describes the contributions to *y* of the cells in *S*_*i-*1_. This is the origin of *y* at the time point *t*_*i-*1_. Similarly, the row *r*(*y,*) of the transition map from *t*_*i*_ to *t*_*i*+1_ describes the probabilities *y* would transition to cells in *S*_*i*+1_. These are the fates of *y*, i.e. the descendants of *y*.

We can compute the origin of *y* further back in time via matrix multiplication: the contributions to *y* of cells in *S*_*i-*2_ are given by a column of the matrix

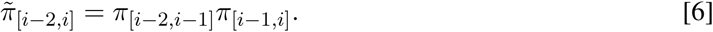

This matrix 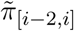 represents the inferred transport from time point *t*_*i-*2_ to *t*_*i*_, we decorate it with a tilde to distinguish it from the maps we compute directly from adjacent time points. Note that we could, in principle, directly compute the transport between any non-consecutive pairs of time points *S*_*i*_, *S*_*j*_, but we do not expect the principle of optimal transport to be as reliable over long time gaps (for more on this see Section 4).

Finally, note that we can interpolate expression profiles between pairs of time points by averaging a cell’s expression profile at time *t*_*i*_ with its fated expression profiles at time *t*_*i*+1_.

###### 3.2 Transport maps encode regulatory information

In this section we demonstrate that transport maps encode regulatory information, and we show how to set up a regression to fit a regulatory function to our sequence of transport maps. We assume that a cell’s trajectory is cell-autonomous and, in fact, depends only on its own internal gene expression. We know this is wrong as it ignores paracrine signaling between cells, and we return to discuss models that include cell-cell communication at the end of this section. However, this assumption is powerful because it exposes the time-dependence of the stochastic process P_*t*_ as arising from pushing an initial measure through a differential equation:

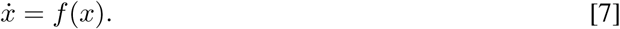

**Figure A3.**
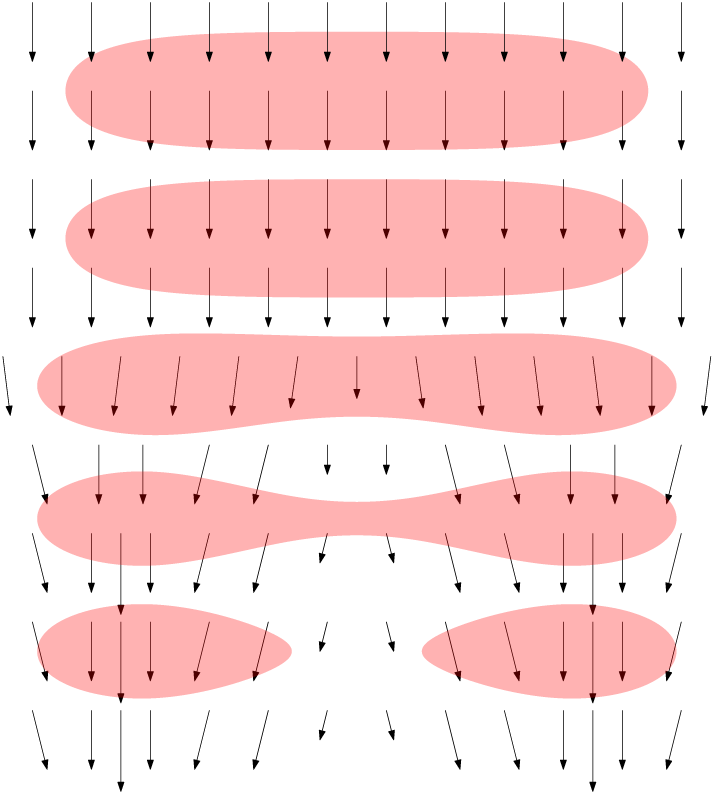
A regulatory vector field gives rise to a time-dependent probability distribution.

Here *f* is a vector field that prescribes the flow of a particle *x* (see **fig. A3** for a cartoon illustration of a distribution flowing according to a vector field). Our biological motivation for estimating such a function *f* is that it encodes information about the regulatory networks that create the equations of motion in gene-expression space.

We propose to set up a regression to learn a regulatory function *f* that models the fate of a cell at time *t*_*i*+1_ as a function of its expression profile at time *t*_*i*_. For motivation that the transport maps might contain information about the underlying regulatory dynamics, we appeal to a classical theorem establishing a dynamical formulation of optimal transport.

**Theorem 1** (Benamou and Brenier, 2001). *The optimal objective value of the transport problem* [1] *is equal to the optimal objective value of the following optimization problem:*

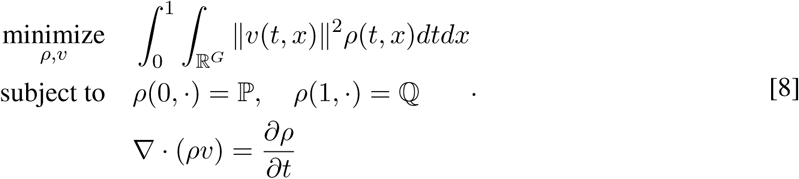

In this theorem, *v* is a vector-valued velocity field that advects^4^ the distribution *ρ* from P to Q, and the objective value to be minimized is the kinetic energy of the flow (mass × squared velocity). Intuitively, the theorem shows that a transport map *π* can be seen as a point-to-point summary of a least-action continuous time flow, according to an unknown velocity field. While the optimization problem [8] can be reformulated as a convex optimization problem, and modified to allow for variable growth rates, it is inherently infinite dimensional and therefore difficult to solve numerically.

We therefore propose a tractable approach to learn a static regulatory function *f* from our sequence of transport maps. Our approach involves sampling pairs of points using the couplings from optimal transport, and solving a regression to learn a regulatory function that predicts the fate of a cell at time *t*_*i*+1_ as a function of its expression profile at time *t*_*i*_:

###### Regulatory network regression

For each pair of time points *t*_*i*_*, t*_*i*+1_, we consider the pair of random variables 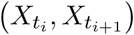 jointly distributed according to 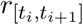, (which we obtained from the transport map 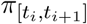 by removing the effect of proliferation as in equation [3]). We set up the following optimization problem over regulatory functions *f*:

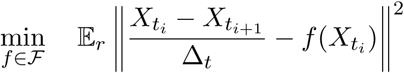

Here *ℱ* specifies a parametric function class to optimize over^5^.

###### Cell non-autonomous processes

We conclude our treatment of gene regulatory networks by discussing an approach to cell-cell communication. Note that the gradient flow [8] only makes sense for cell autonomous processes. Otherwise, the rate of change in expression *x* is not just a function of a cell’s own expression vector *x*(*t*), but also of other expression vectors from other cells. We can accommodate cell non-autonomous processes by allowing *f* to also depend on the full distribution ℙ_*t*_:

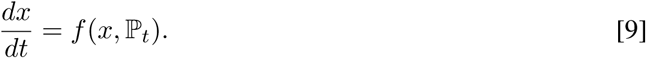

Concretely, we could allow *f* to depend on the mean expression levels of specific genes (expressed by any cell) encoding, for example, secreted factors or direct protein measurements of the factors themselves.

##### 4. Extension to continuous time

In this section we discuss how our method could be improved by going beyond pairs of time points to track the continuous evolution of ℙ_*t*_. We begin by pointing out a peculiar behavior of our method: whenever we have a time point with few sampled cells, our method is forced through an information bottleneck. As an extreme example – suppose we had a time point with only one cell. Everything would transition through that single cell, which is absurd! In this extreme case, we would be better off ignoring the time point. We therefore propose a smoothed approach that shares information between time slices and gracefully improves as data is added.

Our continuous-time formulation is based on locally-weighted averaging, an elementary interpolation technique. Recall that given noisy function evaluations *y*_*i*_ ≈ *f* (*x*_*i*_), one can interpolate *f* by averaging the *y*_*i*_ for all *x*_*i*_ close to a point of interest *x*:

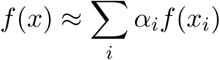

where *α*_*i*_ are weights that give more influence to nearby points (for example 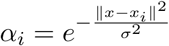.

In our setup, we seek to interpolate a distribution-valued function P_*t*_ from the collections of i.i.d. samples *S*_1_*, …, S*_*T*_. We can interpolate a distribution-valued function by computing the *barycenter* (or centroid) of nearby time points with respect to the optimal transport metric. The *transport barycenter* of a collection of distributions ℙ_1_*, …,* ℙ_*T*_ is a distribution Q that solves the minimization problem

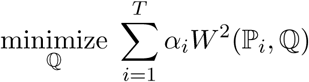

where *W* (ℙ, ℚ) denotes the transport distance (or Wasserstein distance) between P and ℚ. The transport distance is defined by the optimal value of the transport problem [1]. The weights *α*_*i*_ can be chosen to interpolate about time point *t* by setting, for example,

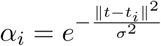

where *∑* is a constant that determines the time-scale for temporal smoothing. Solving for the optimal ℚ is an infinite dimensional optimization problem, but when the support of ℚ is specified ahead of time there are fast algorithms based on entropic regularization (S3).

To apply this in our setting, we modify the transport barycenter to accommodate variable growth rates. We define the gene-expression barycenter of our distributions 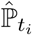 to be the distribution ℚ that solves the minimization problem

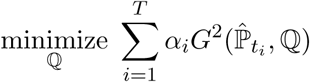

where *G*(ℙ, ℚ) denotes our modified transport distance from equation [5]. To solve this optimization problem, we can fix the support of ℚ to the samples observed at all time points 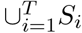*S*_*i*_. Then we can apply the scaling algorithm for unbalanced barycenters due to Chizat et al. (S2).

However, fixing the support of the barycenter ahead of time may not be completely satisfactory, and this motivates further research in the computation of transport barycenters: can we design an algorithm to solve for the barycenter ℚ without fixing the support in advance? Is there a dynamic formulation for barycenters analogous to the Brenier Benamou formula of Theorem 1, and can we leverage it to better learn gene regulatory networks?

Finally, we conclude this section with the observation that this continuous-time approach could provide a principled approach to sequential experimental design. We can identify optimal time points for further data collection by examining the loss function (fit of barycenter) across time, and adding data where the fit is poor. Moreover, we could also use this continuous time approach to test the principle of optimal transport by withholding some time points and testing the quality of the interpolation against the held-out truth. However, a true validation of the principle of optimal transport will require some form of experimental lineage tracing.

### II. Appendix S2

#### WADDINGTON-OT: Concepts and Implementation

In this appendix we describe WADDINGTON-OT: our method for visualizing developmental landscapes and computing developmental trajectories. We begin with an overview in section 1, and we then describe the specific details in sections 2-8.

##### 1. Overview

Recall that we defined in Appendix S1 a notion of the ancestors and descendants of specific subgroups of cells in a population evolving according to a *developmental process*. Moreover, we showed that optimal transport can be used to estimate the temporal evolution of a developmental process from a time-series of scRNA-Seq expression profiles: we compute a transport map between each pair of time points using convex optimization. We also showed how to infer the ancestors and descendants of any group of cells in the dataset, and finally we showed how to fit a regulatory model to the transport maps.

In this appendix we show how to

- compute developmental trajectories
- fit regulatory models to describe their temporal flow
- visualize the developmental landscape.

We also show how to compute clusters of cells and modules of genes, which are helpful tools for exploratory data analysis. Moreover, these cell clusters can be used as subgroups to define ancestor distributions and descendant distributions.

###### Input data

The input to our entire suite of methods is a temporal sequence of single cell gene expression matrices, prepared as described in section 2 below.

###### Computing developmental trajectories and fitting regulatory models

Given a subpopulation of cells, we refer to the sequence of ancestors coming before it and descendants coming after it as its *developmental trajectory*. We describe in section 5 how we compute developmental trajectories in our iPS reprogramming dataset. Briefly, we calculate transport maps between consecutive time points, with cells allowed to grow according to a gene-expression signature of cell proliferation. From these transport maps, the forward or backward transport probabilities can be calculated between any two classes of cells at any time points. For example, we can take successfully reprogrammed cells at day 16 and use back-propagation to infer the distribution over their precursors at day 12. We can then propagate this back to day 11, and so on to obtain the ancestor distributions at all previous time points. This is the developmental trajectory to iPS cells. We can then easily plot trends in gene expression over time (as shown in **Fig. 5** in the main text).

We then fit a regulatory model to the transport maps, as described in section 6. Transcription factors (TFs) that appear to play important roles along trajectories to key destinations are identified by two approaches. The first approach involves constructing a global regulatory model. Pairs of cells at consecutive time points are sampled according to their transport probabilities; expression levels of TFs in the cell at time *t* are used to predict expression levels of all non-TFs in the paired cell at time *t* + 1, under the assumption that the regulatory rules are constant across cells and time points. (TFs are excluded from the predicted set to avoid cases of spurious self-regulation). The second approach involves enrichment analysis. TFs are identified based on enrichment in cells at an earlier time point with a high probability (*>* 80%) of transitioning to a given fate vs. those with a low probability (*<* 20%).

We performed extensive sensitivity analysis to show that key biological results for the reprogramming data are largely robust to the details of the formulation.

###### Visualizing the developmental landscape

To visualize the developmental landscape, we first reduce the dimensionality of the data with diffusion components, and then embed the data in two dimensions with force-directed graph visualization (as described in section 4). While alternative visualization methods, such as t-distributed Stochastic Neighbor Embedding (t-SNE), are well suited for identifying clusters, they do not preserve global structures relevant to studying trajectories across a time course (**fig. S2B,C**). FLE better reflects global structures by including repulsive forces between dissimilar points. In particular, these repulsive forces seem to do a good job of splaying out the spikes present in the diffusion map embedding (**Fig. 3** and **fig. S2A**).

###### Cell clusters and gene modules

Our suite of methods also includes tools for clustering cells (section 7) and identifying modules of correlated genes (section 8).

We cluster cells based on their gene-expression profiles, after performing two rounds of dimensionality reduction to increase statistical power in subsequent analyses. For the reprogramming data, the analysis defined 33 cell clusters. These cell clusters can be used as starting subpopulations to compute developmental trajectories (see Table S4).

Genes are partitioned into sets (*gene modules*) based on correlated expression across the entire dataset, using an approach that searches for tightly connected communities of genes. For the reprogramming data, the analysis partitioned the 16,339 detected genes into 44 gene modules, which were then analyzed for enrichment of gene sets (*signatures*) related to specific pathways, cell types, and conditions (**fig. S3**, **table S2**). Based on the expression profiles in each cell, we calculated signature scores (defined by curated gene sets) for relevant features including MEF identity, pluripotency, proliferation, apoptosis, senescence, X-reactivation, neural identity, placental identity and genomic copy-number variation (CNV). See Appendix S5 for a precise definition of each signature.

##### 2. Preparation of expression matrices

In this section we describe the process by which we compute an expression matrix from scRNA-Seq data. At a high level, we aligned sequenced reads to obtain a matrix *U* of UMI counts, with a row for each gene and a column for each cell. To reduce variation due to fluctuations in the total number of transcripts per cell, we divide the UMI vector for each cell by the total number of transcripts in that cell. Thus we define the expression matrix *E* in terms of the UMI matrix *U* via:

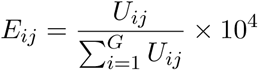

In our subsequent analysis, we make use of two variance-stabilizing transforms of the expression matrix *E*. In particular, we define

1. 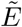 to be the log-normalized expression matrix. The entries of 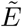 are obtained via

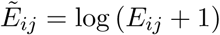
2. Ē to be the truncated expression matrix. The entries of 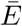are obtained by capping the entries of *E* at the 99.5% quantile.

In the sections below, we denote generic expression profiles as *x* or *y* (these are columns of *E*). Similarly, we denote transformed expression profiles by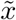 or 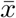 (these are columns of 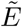 or Ē).

###### Sequencing and alignment of reads

All samples were sequenced to an average depth of 120 million paired-end reads per sample (see SI-3), with 98 bp on the first read and 10 bp on the second read. The 98 bp reads were aligned to the UCSC mm10 transcriptome (using RNA-Seq genome-aligner STAR), and a matrix of UMI counts was obtained using CELLRANGER from the 10X Genomics pipeline (v1.1.0) with default parameters (S4). Quality control metrics about barcoding and sequencing such as the estimated number of cells per collection and the median number of genes detected across cells are summarized in Table S1. To estimate expression of exogenous OKSM factors from OKSM cassette, we extracted RBGpA sequence (839 bp) from the OKSM cassette FASTA file, and generated a reference using the function `mkref` from the CELLRANGER pipeline.

###### Filtering expression matrix

All cells with less than 1000 UMIs detected per cell in total and all genes that were expressed in less than 50 cells were discarded, leaving 65,781 cells and *G* = 16, 339 genes for further analysis. The elements of expression matrix were normalized by dividing UMI count by the total UMI counts detected in a cell and multiplied by 10,000 i.e. expression level is reported as transcripts per 10,000 reads.

##### 3. Dimensionality reduction with diffusion components

Much of our subsequent analysis relies on dimensionality reduction to increase robustness. As a first step towards dimensionality reduction, we remove genes that do not show significant variation (see below for details). We denote the resulting variable-gene expression matrix by *E*_var_.

In our second round of dimensionality reduction, we employ a non-linear mapping called the Laplacian embedding, or diffusion component embedding (S5–S10). While principal components analysis (PCA) is a traditional approach to reduce dimensionality, it is only really appropriate for preserving linear structures. To accommodate nonlinear shapes in high-dimensional gene expression space, we used the diffusion components which are a generalization of principal components (S8).

The diffusion components are defined in terms of a similarity function *k*: ℝ^*G*^×ℝ^*G*^→ [0*,∞*). For a pair (*x, y*) of *G*-dimensional gene-expression profiles, the similarity function—or *kernel* function—*k*(*x, y*) measures the similarity between *x* and *y*. We use the Gaussian kernel function:

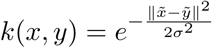

where *x* and *y* are log-transformed expression profiles (i.e. columns of 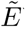).

The diffusion components are defined as the top eigenvectors of a certain matrix constructed by evaluating the kernel function for all pairs of expression profiles *x*_1_*, …, x*_*N*_. Specifically, we form the kernel matrix *K* with entries

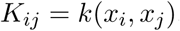

and then we form the Laplacian matrix *L* by multiplying *K* on the left and on the right by *D*,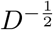 where *D* is a diagonal matrix with entries

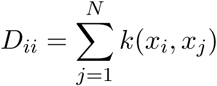

The Laplacian matrix *L* is given by

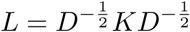

The diffusion components are the eigenvectors *v*_1_*, …, v*_*N*_ of *L*, sorted by eigenvalue. We embed the data in *d* dimensional diffusion component space by selecting the top *d* diffusion components *v*_1_*, …, v*_*d*_, and sending data point *x*_*i*_ to the vector obtained by selecting the *i*th entry of *v*_1_*, …, v*_20_. We denote the diffusion component embedding of an expression profile *x* by ϕ_*d*_(*x*).

We found that the top 20 diffusion components were enriched for gene signatures related to biological processes, and therefore we elected to use the top 20 diffusion components to represent our data (see below for details).

###### Selecting variable genes

We used the function `MeanVarPlot` from the SEURAT package (S11) (v1.4.0.6) to select 2076 variable genes. First, we divided genes into 20 bins based on their average expression levels across all cells. Second, we compute Fano factor of gene expression in each bin and then z-scored. The Fano factor, defined as the variance divided by the mean, is a measure of dispersion. Finally, by thresholding the z-scored dispersion at 0.5, we obtained a set of 2076 variable genes.

###### Computing diffusion components

We ran the R package DESTINY (S8) with the following parameters:

- `sigma=’local’` (This option calculates the diffusion scale parameter of Gaussian kernel separately for every cell based on the distances to its nearest neighbors in gene expression space),
- `k=250` (This specifies the number of nearest neighbors for which the entries of the kernel matrix are evaluated. Others are left as 0. We selected *k* = because it is close to the square root of the total number of cells.)
- `n_eigs=100` (This specifies the number of diffusion components to compute).

Finally, we examined the enrichment of the top 100 diffusion components for biological signatures, as described below. We found that the top 20 diffusion components were enriched for biological terms related to development (e.g. diffusion component 20 is related to placenta development). We therefore elected to use the top 20 diffusion components.

We computed three separate sets of diffusion components: for 2i and serum together, and also for each separately.

###### Selection of diffusion components by biological enrichment

We analyzed the biologicalprocess enrichment of the top 100 diffusion components to select the dimensionality of the embedding ϕ_*d*_. For each diffusion component, we identified the genes whose expression levels were correlated with the numerical values of the diffusion component across cells. We took the genes with absolute correlation coefficient exceeding 0.2. This gave us a set of genes for each diffusion component. We then performed functional enrichment analysis on these gene sets using the `findGO.pl` program from HOMER (S12) with Benjamini and Hochberg correction for multiple hypothesis testing (retaining terms at adjusted *p*-value *<* 0.05). We found that the top 20 diffusion components contained information relevant to developmental processes, and therefore we elected to use the top 20 diffusion components in our subsequent analysis.

##### 4. Visualization: force-directed layout embedding

In this section we introduce our two dimensional visualization technique based on force-directed layout embedding (FLE) (S13, S14). FLE is large-scale graph visualization tool which simulates the evolution of a physical system in which connected nodes experience attractive forces, but unconnected nodes experience repulsive forces. Initial FLE algorithms used simple electrostatic and spring forces, but modern FLE algorithms allow for more elaborate interactions that can depend on the degree of nodes or include gravity terms that attract all nodes to the center (this is especially important for disconnected graphs, which would otherwise fly apart). Starting from a random initial position of vertices, the network of nodes evolves in such a manner that at any iteration a new position of vertices is computed from the net forces acting on them. This procedure resembles the Molecular Dynamics simulation method for studying dynamics of atoms in molecules.

We apply the FLE to visualize the nearest neighbor graph generated from our data. Our intuition behind applying such a visualization procedure, with long-range repulsive forces between distant components, was that it would splay out the spikes visible in different diffusion components (see **Fig. 3** and **fig S2A**).

###### Implementation

First, we reduce from 16, 000 dimensions to 20 dimensions by computing a 20 dimensional diffusion component embedding of the dataset ϕ_20_(*x*_1_)*, …* ϕ_20_(*x*_*N*_).

Second, for each cell we compute its 20 nearest neighbors in 20-dimensional diffusion component space to produce a nearest neighbor graph. For this step, we used the fast k-NN algorithm from the R package FNN (S15).

Finally, we compute the force-directed layout on the k-NN graph using the ForceAtlas2 algorithm. It took around 1 h to reach steady-state on a single machine using the `ForceAtlas2` algorithm (S14) from the Gephi Toolkit (v0.9.1) (S13). This algorithm has been previously used to visualize highdimensional single-cell mass cytometry data (CyTOF) of measured levels of marker proteins (up to 50 markers per single cell) (S16–S18).

##### 5. Computing trajectories with optimal transport

Recall that for any pair of time points we compute a transport plan that minimizes the expected cost of redistributing mass, subject to constraints involving a proliferation score (see Appendix 1 for a precise statement of the optimization problem). To compute these transport matrices, we need to specify a cost function, a proliferation function, and numerical values for the regularization parameters.

###### Cost functions

We tried several different cost functions based on squared Euclidean distance in different input spaces. Specifically, for cells with expression profiles *x* and *y*, given by two columns of the expression matrix *E*, we specify a cost function *c*(*x, y*)

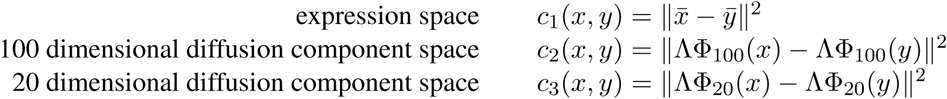

The bar above 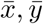 denotes that we apply the truncation transform from section 2, and ϕ_*d*_ is the Laplacian embedding from section 3. Note that ϕ_*d*_ has the log transform 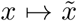 built-in. In the equations above, λ is a diagonal matrix containing the eigenvalues of the Laplacian matrix, raised to the power 8. Hence *c*_2_ and *c*_3_ are both truncated versions of the diffusion distance *D*_4_(*x, y*) from (S5).

We used the cost function *c*_3_ to report the numerical values in the main text, and we computed separate transport maps for 2i and serum. Note that all the cost functions *c*_1_*, c*_2_*, c*_3_ give largely similar results.

###### Proliferation function

We estimate the *relative* growth rate for every cell using the proliferation signature displayed in **Fig. 3D** in the main text. To transform the proliferation score into an estimate of the growth rate (in doublings per day), we first observed that the proliferation score is bimodally distributed over the dataset. We transformed the proliferation score so that the two modes were mapped to a growth ratio of 2.5 per day (this means that over 1 day, a cell in the more proliferative group is expected to produce 2.5 times as many offspring as a cell in the non-proliferative group). However, note that we allow for some laxity in the prescribed growth rate (see supplemental figure on input vs implied proliferation).

###### Regularization parameters

We employed the following strategy to select the regularization parameters *λ* and `∈`. The *entropy* parameter `∈` controls the entropy of the transport map. An extremely large entropy parameter will give a maximally entropic transport map, and an extremely small entropy parameter will give a nearly deterministic transport map (but could also lead to numerical instability in the algorithm). We adjusted the entropy parameter until each cell transitions to between 10 and 50 percent of cells in the next time point, as measured by the Shannon diversity of the rows of the transport map.

The regularization parameter *λ* controls the fidelity of the constraints: as *λ* gets larger, the constraints become more stringent. We selected *λ* so that the marginals of the transport map are 95% correlated with the prescribed proliferation score.

###### Implementation

We implemented the scaling algorithm for unbalanced transport (S2) to compute optimal transport maps. This algorithm performs gradient ascent steps on the dual optimization problem. Because of the entropic regularization, these gradient ascent steps can be performed via diagonal matrix scalings. We implemented versions of the solver in both R and Python.

###### Experiments

We performed computational experiments to evaluate the stability of our results to choice of cost function, regularization parameters, and subsampling the dataset.

- We compared the cluster-to-cluster origin and fate tables for the different cost functions listed above, and we found reassuringly consistent results. Moreover, the transport probabilities mentioned in the main text are all robust to choice of cost function.
- We performed a bootstrap analysis on a batch of 100 subsamples consisting of 50% of the data from each time point. The variance in the cluster-to-cluster origin and fate tables is extremely small (see Table S4).
- As an additional validation, we modified an existing trajectory finding technique, Wishbone (S10)— based on shortest paths in k-NN graphs—to include information about time and proliferation. This gives trajectories (shown in **fig. ??**) whose overall shape agrees with the transports displayed in **Fig. 4A**. We describe the implementation of our modified k-NN approach in *Appendix S4*.

##### 6. Learning gene regulatory networks

Recall that in Appendix S1 we showed how to set up an optimization problem to solve for a regulatory function that fits the transport maps.

In order to make this concrete, we need to specify a function class over which to optimize (see section 3.2 of Appendix S1). We consider a rectified-linear function class defined in terms of a specific generalized logistic function *𝓁*: ℝ *1→* ℝ:

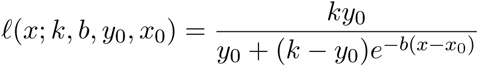

where *k, b, y*_0_*, x*_0_ *∈* ℝ are parameters of the generalized logistic function *𝓁*(*x*).

We define a function class 𝔽 consisting of functions *f*: ℝ^*G*^ *→* ℝ^*G*^ of the form

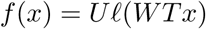

where 𝓁 is applied entry-wise to the vector *WZx ∈* ℝ^*M*^ to obtain a vector that we multiply against *U ∈* ℝ^*G×M*^. Here 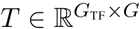 denotes a projection operator that selects only the coordinates of *x* that are transcription factors, and *G*TF is the number of transcription factors.

We set up the following optimization over matrices *U ∈* ℝ^*G×M*^ and 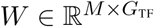:

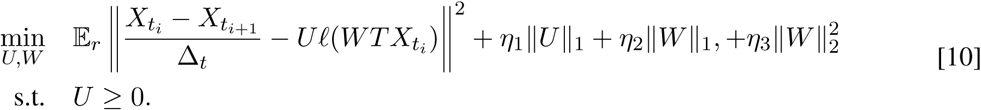

where 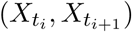 is a pair of random variables distributed according to the normalized transport map *r* (as in section 3.2 of Appendix S1), and ∥*U*∥_1_ denotes the sparsity-promoting *𝓁*_1_ norm of *U*, viewed as a vector (that is, the sum of the absolute value of the entries of *U*). Each rank one component (row of *U* or column of *W*) gives us a group of genes controlled by a set of transcription factors. The regularization parameters *η*_1_ and *η*_2_ control the sparsity level (i.e. number of genes in these groups).

###### Implementation

We designed a stochastic gradient descent algorithm to solve [10]. Over a sequence of epochs, the algorithm samples batches of points 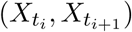 from the transport maps, computes the gradient of the loss, and updates the optimization variables *U* and *W*. The batch sizes are determined by the Shannon diversity of the transport maps: for each pair of consecutive time points, we compute the Shannon diversity *S* of the transport map, then randomly sample max(*S ×* 10^−5^, 10) pairs of points to add to the batch. We run for a total of 10, 000 epochs.

We implemented this algorithm in Python.

##### 7. Clustering cells

We cluster cells using the Louvain-Jaccard community detection algorithm (S19–S21) in 20 dimensional diffusion component space. This algorithm maximizes the *Louvain modularity* — a value between *-*1 and 1 that measures the density of links inside communities compared to links between communities.

As a first step, we compute the 20-nearest neighbor graph in 20 dimensional diffusion component space (computed on cells from both 2i and serum). We then weight the edges in this graph by the Jaccard similarity coefficient. We partition the resulting graph into clusters using the Louvain community detection algorithm (S19) implemented in the function `multilevel.community` from the R package IGRAPH (1.0.1) (S22). The default parameters for automatically selecting the number of clusters gave us 33 clusters, displayed in **Fig. 3D** in the main text.

##### 8. Gene correlation modules reveal biological signatures

In this section we describe our technique for identifying modules of correlated genes, with the goal of revealing coherent biological processes.

Our procedure consists of two steps. In the first step, we use the Graphical Lasso (S23) to compute a regularized estimate of the covariance matrix for our 66,000 expression profiles. The Graphical Lasso fits a covariance matrix to the data, regularized so that the inverse of the covariance matrix is *sparse* (i.e. has only a few non-zeros). The motivation for selecting a sparse inverse covariance is based on the fact that if a collection of observations have a multivariate Gaussian distribution with mean *μ* and covariance ∑, then the zero pattern of ∑^−1^ completely specifies the conditional independence structure of the observations:

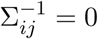 *⇔* variables *i* and *j* are conditionally independent given the other variables. Let θ = ∑^−1^ and let *S* denote the empirical covariance for our expression profiles

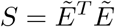

The Graphical Lasso maximizes the Gaussian log likelihood:

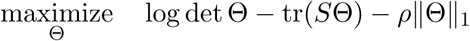

Here ∥Θ ∥_1_ is a regularization term that promotes sparse solutions. The optimal Θ is a (regularized) maximum-likelihood estimate of the inverse covariance matrix ∑^−1^ for a Gaussian ensemble.

We identify gene modules as tightly knit communities in the network specified by Θ (see below). Based on these gene modules, we then identified gene signatures related to specific pathways, cell types, and conditions. We did this by functional enrichment analysis (see below). The gene modules are displayed in **fig. S3**.

###### Computing gene modules

We used the `glasso` package (S23) to solve the graphical lasso optimization problem. We tune the regularization parameter *ρ* to achieve a desirable sparsity level for Θ. In particular, we select a value of *ρ* that gave around 10, 000 total genes (i.e. 10, 000 non-zero rows and columns of Θ).

Viewing θ as an adjacency matrix defining a network of genes, we partitioned the network using with the Infomap community detection algorithm (S24) from the R package IGRAPH (v1.1.0) (S22), retaining modules that contain more than 10 genes. This yields 44 gene modules, each consisting of a set of genes. The modules are visualized in **fig. S3**.

###### Functional enrichments

We performed functional enrichment analysis on the gene sets defined by the modules using the `findGO.pl` program from the HOMER suite (Hypergeometric Optimization of Motif Enrichment, version: 4.9.1) (S12) with Benjamini and Hochberg correction for multiple hypothesis testing (retaining terms at adjusted *p*-value *<* 0.05). All genes that passed quality-control filters were used as a background set.

This yielded a set of biological signatures related to each module. See Appendix S5 for a precise definition of the relevant signatures (including MEF identity, pluripotency, proliferation, apoptosis, senescence, X-reactivation, neural identity and placental identity).

###### Computing scores from gene sets

Given a set of genes (coming from a gene module or biological signature), we score cells based on their gene expression. In particular, for a given cell we compute the z-score for each gene in the set. We then truncate these z-scores at 5 or 5, and define the signature of the cell to be the mean z-score over all genes in the gene set. The scores for the gene modules are visualized in **fig. S3** and the scores for the biological signatures are visualized in **Fig. 3**.

### III. Appendix S3

#### Experimental Methods

In this appendix we describe our experimental methods.

##### 1. Derivation of MEFs

Mouse embryonic fibroblasts (MEFs) were derived from E13.5 embryos with a mixed B6;129 background. The cell line used in this study was homozygous for ROSA26-M2rtTA, homozygous for a polycistronic cassette carrying *Pou5f1*, *Klf4*, *Sox2*, and *Myc* at the *Col1a1* locus (18), and homozygous for an EGFP reporter under the control of the *Pou5f1* promoter. Briefly, MEFs were isolated from E13.5 embryos resulting from timed-matings by removing the head, limbs, and internal organs under a dissecting microscope. The remaining tissue was finely minced using scalpels and dissociated by incubation at 37°C for 10 minutes in trypsin-EDTA (Thermo Fisher Scientific). Dissociated cells were then plated in MEF medium containing DMEM (Thermo Fisher Scientific), supplemented with 10% fetal bovine serum (GE Healthcare Life Sciences), non-essential amino acids (Thermo Fisher Scientific), and GlutaMAX (Thermo Fisher Scientific). MEFs were cultured at 37°C and 4% CO_2_ and passaged until confluent. All procedures, including maintenance of animals, were performed according to a mouse protocol (2006N000104) approved by the MGH Subcommittee on Research Animal Care.

##### 2. Reprogramming assay

For the reprogramming assay, 20,000 low passage MEFs (no greater than 3-4 passages from isolation) were seeded in a 6-well plate. These cells were cultured at 37°C and 5% CO_2_ in reprogramming medium containing KnockOut DMEM (GIBCO), 10% knockout serum replacement (KSR, GIBCO), 10% fetal bovine serum (FBS, GIBCO), 1% GlutaMAX (Invitrogen), 1% nonessential amino acids (NEAA, Invitrogen), 0.055 mM 2-mercaptoethanol (Sigma), 1% penicillin-streptomycin (Invitrogen) and 1,000 U/ml leukemia inhibitory factor (LIF, Millipore). Day 0 medium was supplemented with 2*μ*g/mL doxycycline Phase-1(Dox) to induce the polycistronic OKSM expression cassette. Medium was refreshed every other day. At day 8, doxycycline was withdrawn, and cells were transferred to either serum-free 2i medium containing 3 *μ*M CHIR99021, 1 *μ*M PD0325901, and LIF (Phase-2(2i)) (25) or maintained in reprogramming medium (Phase-2(serum)). Fresh medium was added every other day until the final time point on day 16. Oct4-EGFP positive iPSC colonies should start to appear on day 10, indicative of successful reprogramming of the endogenous *Oct4* locus.

##### 3. Sample collection

We collected a total of 66,000 cells from twelve time points over a period of 16 days in two different culture conditions. Single or duplicate samples were collected at day 0 (before and after Dox addition), 2, 4, 6, and 8 in Phase-1(Dox); day 9, 10, 11, 12, 16 in Phase-2(2i); and day 10, 12, 16 in Phase-2(serum). Cells were also collected from established iPSCs cell lines reprogrammed from the same MEFs, maintained either in Phase-2(2i) conditions or in Phase-2(serum) medium. For all time points, selected wells were trypsinized for 5 mins followed by inactivation of trypsin by addition of MEF medium. Cells were subsequently spun down and washed with 1X PBS supplemented with 0.1% bovine serum albumin. The cells were then passed through a 40 micron filter to remove cell debris and large clumps. Cell count was determined using Neubauer chamber hemocytometer to a final concentration of 1000 cells/*μ*l.

##### 4. Single-cell RNA sequencing

Single-cell RNA-Seq libraries were generated from each time point using the 10X Genomics Chromium Controller Instrument (10X Genomics, Pleasanton, CA) and Chromium^TM^ Single Cell 3’ Reagent Kits v1 (PN-120230, PN-120231, PN-120232) according to manufacturer’s instructions. Reverse transcription and sample indexing was performed using the C1000 Touch Thermal cycler with 96-Deep Well Reaction Module. Briefly, the suspended cells were loaded on a Chromium controller Single-Cell Instrument to first generate single-cell Gel Bead-In-Emulsions (GEMs). After breaking the GEMs, the barcoded cDNA was then purified and amplified. The amplified barcoded cDNA was fragmented, Atailed and ligated with adaptors. Finally, PCR amplification was performed to enable sample indexing and enrichment of the 3’ RNA-Seq libraries. The final libraries were quantified using Thermo Fisher Qubit dsDNA HS Assay kit (Q32851) and the fragment size distribution of the libraries were determined using the Agilent 2100 BioAnalyzer High Sensitivity DNA kit (5067-4626). Pooled libraries were then sequenced using Illumina Sequencing By Synthesis (SBS) chemistry.

##### 5. Lentivirus Vector Construction and Particle Production

To test whether transcription factors (TFs) improve late-stage reprogramming efficiency, we generated lentiviral constructs for the top candidates *Zfp42*, and *Obox6*. cDNA for these factors were ordered from Origene (*Zfp42*-MG203929, and *Obox6*-MR215428) were cloned into the FUW Tet-On vector (Addgene, Plasmid #20323) using the Gibson Assembly (NEB, E2611S). Briefly, the cDNA for each TF was amplified and cloned into the backbone generated by removing Oct4 from the FUW-Teto-Oct4 vector. All vectors were verified by Sanger sequencing analysis. For lentivirus production, HEK293T cells were plated at a density of 2.6 × 10^6^ cells/well in a 10cm dish. The cells were transfected with the lentiviral packaging vector and a TF-expressing vector at 70-80% growth confluency using the Fugene HD reagent (Promega E2311) according to the manufacturer’s protocols. At 48 hours after transfection, the viral supernatant was collected, filtered and stored at -80°C for future use.

##### 6. Reprogramming efficiency of secondary MEFS together with individual TFs

We sought to determine the ability of the candidate TFs to augment reprogramming efficiency in secondary MEFs; the use of secondary MEFs for reprogramming overcomes limitations associated with random lentiviral integration events at variable genomic locations. Briefly, secondary MEFs were plated at a concentration of 20,000 cells per well of a 6-well plate. Cells were infected with virus containing *Zfp42*, *Obox6*, or an empty vector and maintained in reprogramming medium as described above. At day 8 after induction, cells were switched to either Phase-2(2i) or Phase-2(serum). On day 16, reprogramming efficiency was quantified by measuring the levels of the EGFP reporter driven by the endogenous *Oct4* promoter. FACS analyses was performed using the Beckman Coulter CytoFLEX S, and the percentage of Oct4-EGFP^+^ cells was determined. Triplicates were used to determine average and standard deviation (**Fig. 6B**).

##### 7. Reprogramming efficiency of primary MEFS with individual TFs and OKSM

In addition to demonstrating the ability of a TF to increase reprogramming efficiency in secondary MEFs, we sought to independently test performance in primary MEFs. To this end, lentiviral particles were generated from four distinct FUW-Teto vectors, containing *Oct4*, *Sox2*, *Klf4*, and *Myc*, previously developed in the Jaenisch lab. MEFs from the background strain

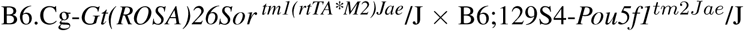

were infected with these lentiviral particles, together with a lentivirus expressing tetracycline-inducible *Zfp42*, *Obox6* or no insert. Infected cells were then induced with 2*μ*g/mL doxycycline in ESC reprogramming medium (day 0). At day 8 after induction, cells were switched to either Phase-2(2i) or Phase-2(serum). On day 16, the number of Oct4-EGFP^+^ colonies were counted using a fluorescence microscope. Triplicates for each condition used to determine average values and standard deviation.

### IV. Appendix S4

#### Performance of Published Methods

In this appendix we review published methods for identifying cellular trajectories from scRNA-Seq data, and we investigate their performance on our dataset. We emphasize in fairness that we applied these methods out-of-the-box, without optimizing their performance for our specific data.

As we note in the main text, these methods were initially designed to extract information about stationary processes (such as the cell cycle or adult stem cell differentiation) in which all stages exist simultaneously. As a result, they neither directly model nor explicitly leverage the temporal information in our developmental time course. We find that they all run into problems in a system such as ours because they ignore time (Sections 1-3), and we explain why in Section 4.

In addition to the above issue about time, these methods often also impose strong structural constraints on the model, such as one-dimensional trajectories and zero-dimensional bifurcation points. Indeed, most papers in the literature represent developmental trajectories with a graph embedded in gene expression space, where nodes represent differentiation events as zero-dimensional points, and edges represent developmental trajectories as one-dimensional paths. There are several fundamental limitations to representing developmental processes mathematically by a graph embedded in gene expression space:

- A graph itself does not encode the *probability* that a cell at a given time will lead to descendants with a particular fate.
- While each node could be annotated with the likelihood of traveling down each outgoing edge, a graph cannot describe continuous changes in fate probabilities. The probability of arriving at a terminal state must change in a discrete, step-wise fashion at each node and must be constant along edges.
- A graph can only describe simple branch points, where there is a clean split between lineages at a well-defined position. Consider a hypothetical situation where at any point en route from *A* to *B*, one can split off and head towards *C*. The set of possible trajectories could fill the interior of a triangle with vertices *ABC*. The bifurcation event is not localized to a zero-dimensional point, represented by the node of a graph.
- A graph can only describe trajectories to finitely many states. In a hypothetical situation where cells differentiate from a single state towards a continuum of states, the set of possible trajectories to the continuum of terminal states might look like a triangle with one vertex representing the origin and the opposite side representing the continuum of terminal states.
- When the graph is restricted to be a tree, cells cannot arrive at a state in more than one way.

##### 1. The diffusion pseudo-time method (DPT)

The diffusion pseudo-time method created by Haghverdi et al. in 2016 (S7) orders cells using an appropriately weighted distance in diffusion component space: the diffusion components *ϕ*_*i*_ are rescaled by 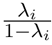, where *λ*_*i*_ is the eigenvalue for the *i*th diffusion component vector, *ϕ*_*i*_. This rescaling is derived by considering a random walk according to a transition matrix *T* that specifies the probability of transitioning from any single cell to another in an infinitesimal amount of time. Therefore, at some level, the DPT method can also deal with time-varying distributions on gene expression space: the time evolution of a probability density *f* (*t*) is described by

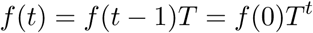

This is of course true in general, but the crux is to specify *T*. Note that in our method, we do this with optimal transport, from one time slice to the next. By contrast, the diffusion pseudotime method (DPT) is designed for stationary processes, and assumes by necessity that there is no information available about time. It simply constructs a transition matrix *T* from a kernel function *k*:

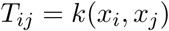

The rescaling by 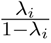 takes into account the effect of summing over walks of varying length, and makes the diffusion distance between *x* and *y* proportional to the accumulated probability of transitioning from *x* to *y*.

The diffusion pseudo-time method orders cells according to the DPT distance from a certain starting cell *x*_0_ (which can be selected automatically). The method identifies branch points as regions where the triangle inequality for the DPT distance is strongly violated (the triangle inequality is most violated at the tip of an off-shoot branch).

Applied to our data, the diffusion pseudo-time does not order cells correctly — it places large numbers of day 9 cells (in the valley) before cells collected on day 2. (We discuss the reason for this in Section 4 below). Figure A4 illustrates the diffusion pseudo-time of cells from the serum condition, starting from a random day 0 cell. Panel A shows the distribution of diffusion pseudo-times for each real time. Each point corresponds to a single cell. The cells are colored by their diffusion pseudo-time. Panel B shows the distribution of diffusion pseudo-time over the force-directed layout embedding. Note that the cells in the valley are almost all closer to day 0 than any of the day 2 cells are. In particular, there are only 50 cells from day 2 with DPT less than 0.21 (horizontal line in A), while there are more than 1000 such cells from days 10−16.

Therefore the ordering computed by diffusion pseudo-time is temporally impossible.

##### 2. Monocle 2

Monocle 2 from X. Qiu et al. (S25, S26) fits a principal graph (S27) to a collection of scRNA-Seq profiles. The algorithm is based on a technique called reversed graph embedding which simultaneously learns a low dimensional embedding of the data and a graphical structure spanning the dataset. The search for a low dimensional embedding is initialized with principal components analysis, and for computational reasons the computational search is restricted to trees using the DDRTree algorithm (S27). As for DPT, the method assumes that time information is not available.

We applied DDRTree (S27) (R package DDRTree, Ver. 0.1.5) to a downsampled expression matrix of variable genes (2076 genes and 30,000 cells), because we could not get the algorithm to run on our entire dataset with our computational resources (The algorithm took 6 days to run on the downsampled dataset on an Intel Xeon CPU, 200 GB of memory).

**Figure A4.**
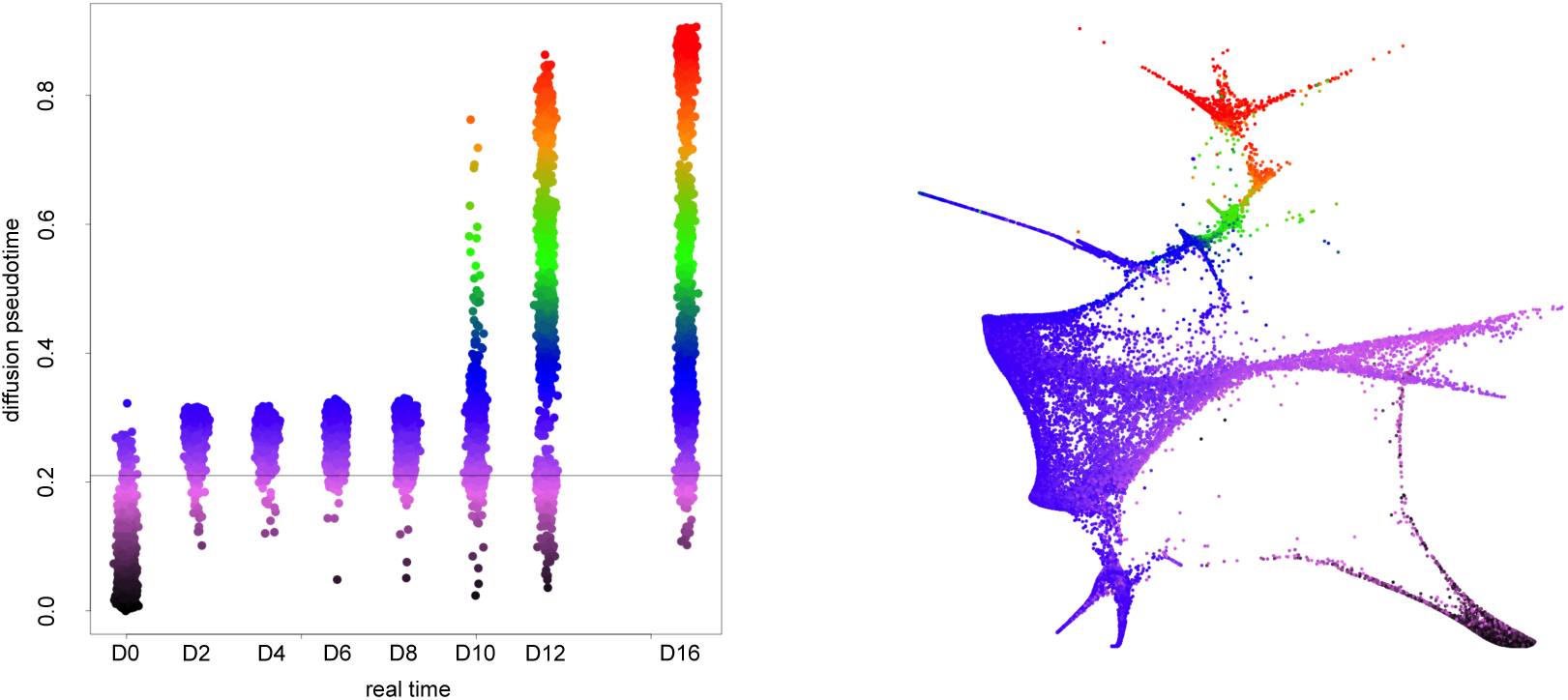
Diffusion pseudo-time distance from day 0. Panel (A) shows a plot of pseudo-time vs real time, for cells grown in serum. The points are colored by pseudo-time, and the real-time values are jittered to show the density of points. Panel (B) shows the force layout embedding visualization, also colored by pseudo-time. The pseudotime ordering erroneously places a large number of day 12 cells in the valley before most day 2 cells.

Various trajectories were, however, clearly incompatible with the time course (fig. A5). For example, the graph has an early bifurcation at day 0, with cells transitioning directly to cells at days 10-16 in the Valley (without passing through cells at intermediate times) and the remaining cells on the trajectory to successful reprogramming.

##### 3. Wishbone

Wishbone, from M. Setty et al. (S10), uses shortest paths in a k-nearest neighbor (kNN) graph constructed in diffusion component space to construct an initial ordering of cells. These paths start from a user-defined cell. Next, the initial ordering is refined by considering all shortest paths to and from a randomly selected population of waypoint cells. Branch-points are identified by clustering the waypoint cells according to the shortest path distance they assign to all nearby cells, using spectral clustering. The final output of the Wishbone program is a branching tree that is fit to the kNN graph in this way.

Applied to our dataset, Wishbone identifies a branching structure which is inconsistent with the correct temporal sequence, as demonstrated in **fig. A6**. Wishbone identifies a branching event in day 0 and partitions all subsequent days into two branches, which we refer to as Branch B and Branch C (day 0 is Branch A, the trunk connected to the root of the tree). Branch C contains all cells from days 2 → 8, and Branch B contains and all cells from days 10 → 16. Since both branches connect directly to Branch A, which itself consists of only day 0 cells, this branching structure is inconsistent with the measured time information. We modify Wishbone to add time information below.

**Figure A5.**
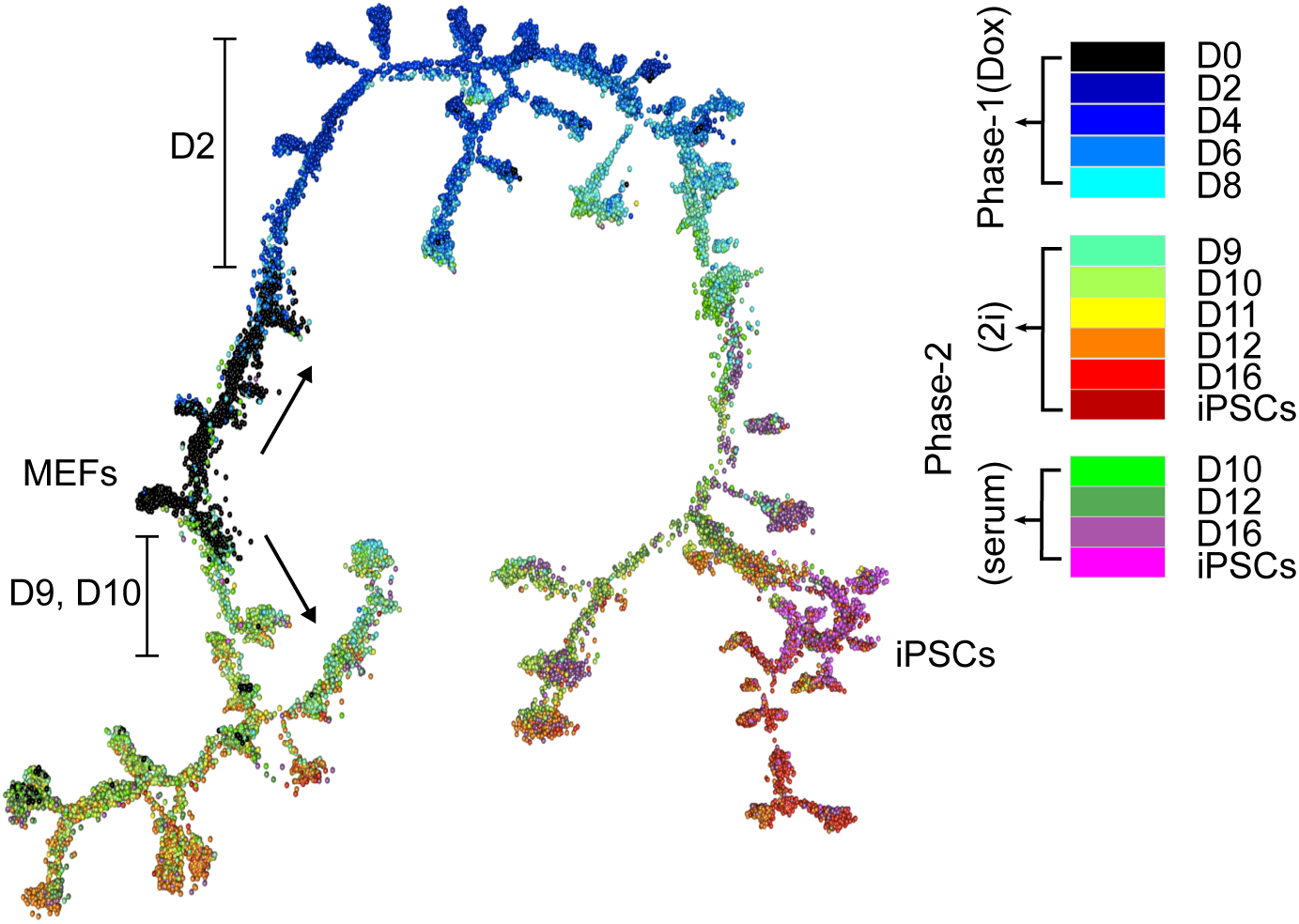
Monocle2 trajectories include an impossible temporal jump directly from day 0 MEFs to day 9 cells in the ”valley”. Cells are colored by real time.

##### 4. What goes wrong without time?

The inability to incorporate time information is especially limiting in situations where cells can revert to an earlier state. Moreover, this problem is magnified for programs designed to represent differentiation processes as trees; they erroneously split such (undirected) loops into distinct branches. This mode of failure, which we observe across all published methods applied to our dataset, is illustrated with a cartoon example in **fig A7**. Panel (a) of **fig A7** shows a schematic example of a bifurcating trajectory where one branch reverts to an earlier state. The colors represent time information (after the bifurcation event, contemporaneous samples are represented by two dots with the same color). Panel (b) shows a trajectory inferred by connecting nearby samples, without using time information (colors are still shown for clarity).

In our biological system, MEF reversion creates the situation illustrated in **fig. A7**. Indeed, as shown above, diffusion pseudo-time (S7) (Section 1), Monocle 2 (S25, S26) (Section 2), and Wishbone (S10) (Section 3) all fail in this way. In section **??** we show how to modify a kNN approach motivated by Wishbone and DPT to incorporate information about time. This fixes the particular issue with MEF reversion. However, as a graph-based method it still faces the other fundamental issues listed at the beginning of this Appendix.

##### 5. TASIC: a method that incorporates time information

While preparing our manuscript, we became aware of TASIC (S28) — a method that is able to incorporate time information and identify branches in single cell expression data. TASIC models developmental processes as emitting expression profiles from a finite number of states. Over the course of time, cells are allowed to hop from one discrete state to another. The measured expression profile observed from any given state is assumed to be Gaussian in expression space, and the temporal duration of the state is assumed to be Poisson.

**Figure A6.**
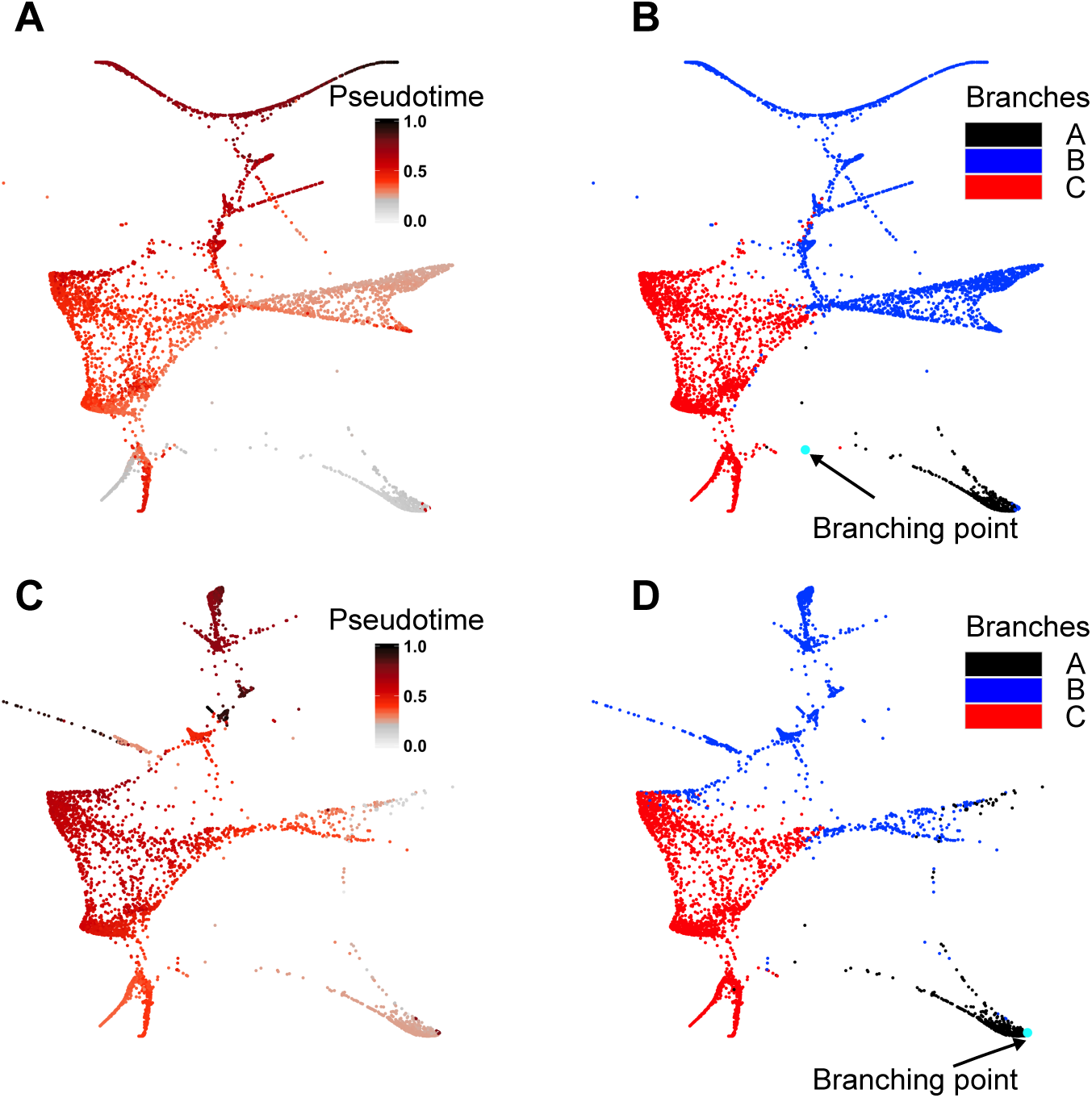
Wishbone identifies a major branching event at day 0 and partitions the cells from days 2 *→* 16 into two groups — one containing cells from day 2 *→* 8 and one containing cells from day 10 *→* 16.

However, there are several challenges in this method.

- The assumption that expression profiles follow a Gaussian distribution is well known to be unrealistic for single cell RNA-Seq data. When there are very few transcripts per cell, the central limit theorem does not guarantee a Gaussian distribution.
- The model only allows cells to occupy finitely many states, and the transitions are computed between states.
- The computational cost of fitting the model is exponential in the number of states considered, making the algorithm impractical with more than a few time points.

We tested TASIC on our dataset (**fig A8**) with the following setting: 2076 variable genes, cells across all timepoints were downsampled to 5400 cells (we consider here only 2i condition). We ran TASIC with the following number of states: 1) 3 states (**fig. A8A**), 2) 5 states (**fig. A8B)**, and 3) 7 states (**fig. A8C**). For each number of states we obtained monolineage trajectories (**fig. A8**, bottom panel).

**Figure A7.**
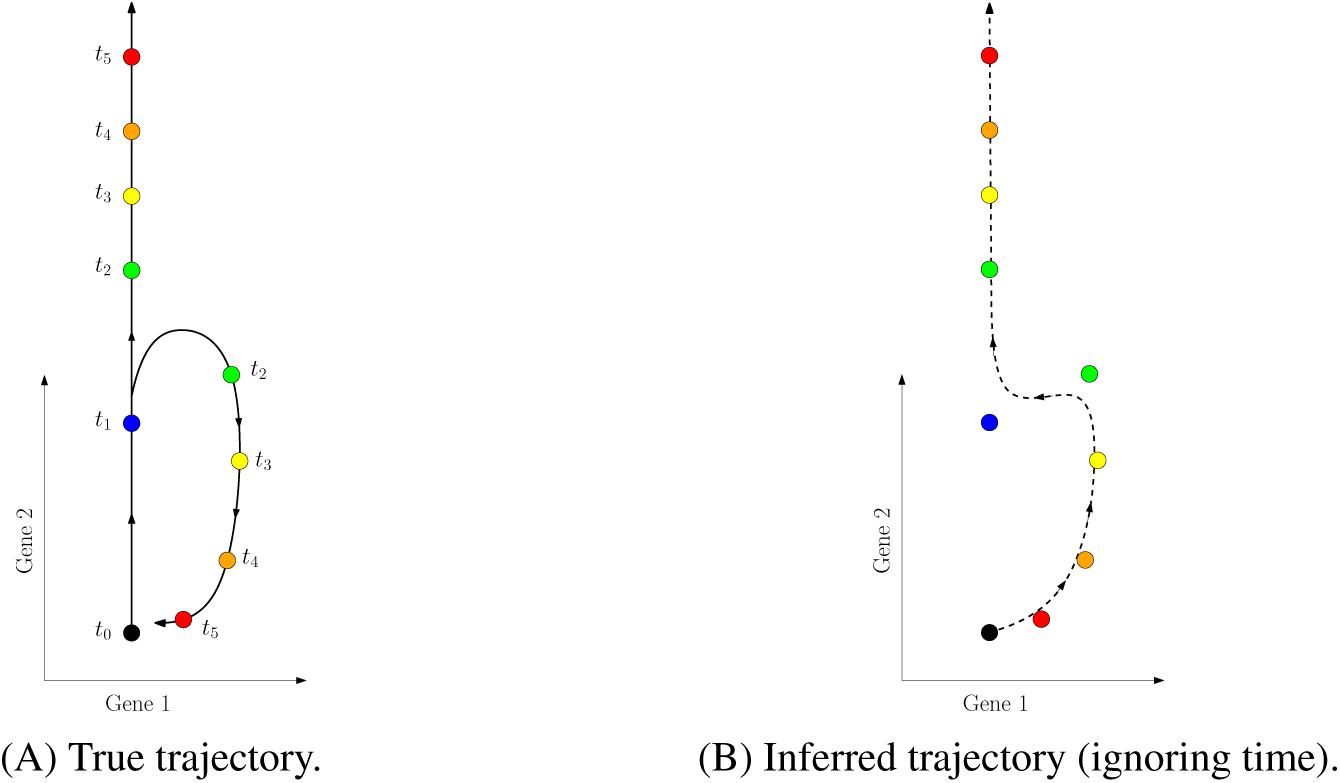
Time information is especially useful when cells can reach the same fate through different routes. (A) A schematic trajectory in two dimensional gene expression space with one bifurcation. After the bifurcation, one branch returns to the starting position. Samples are color coded by time of collection (note that after the bifurcation point, the samples are bimodal and are represented by two dots with the same color). (B) A schematic of a trajectory inferred by connecting nearby samples without using time information (a smoothed minimum spanning tree is drawn).

**Figure A8.**
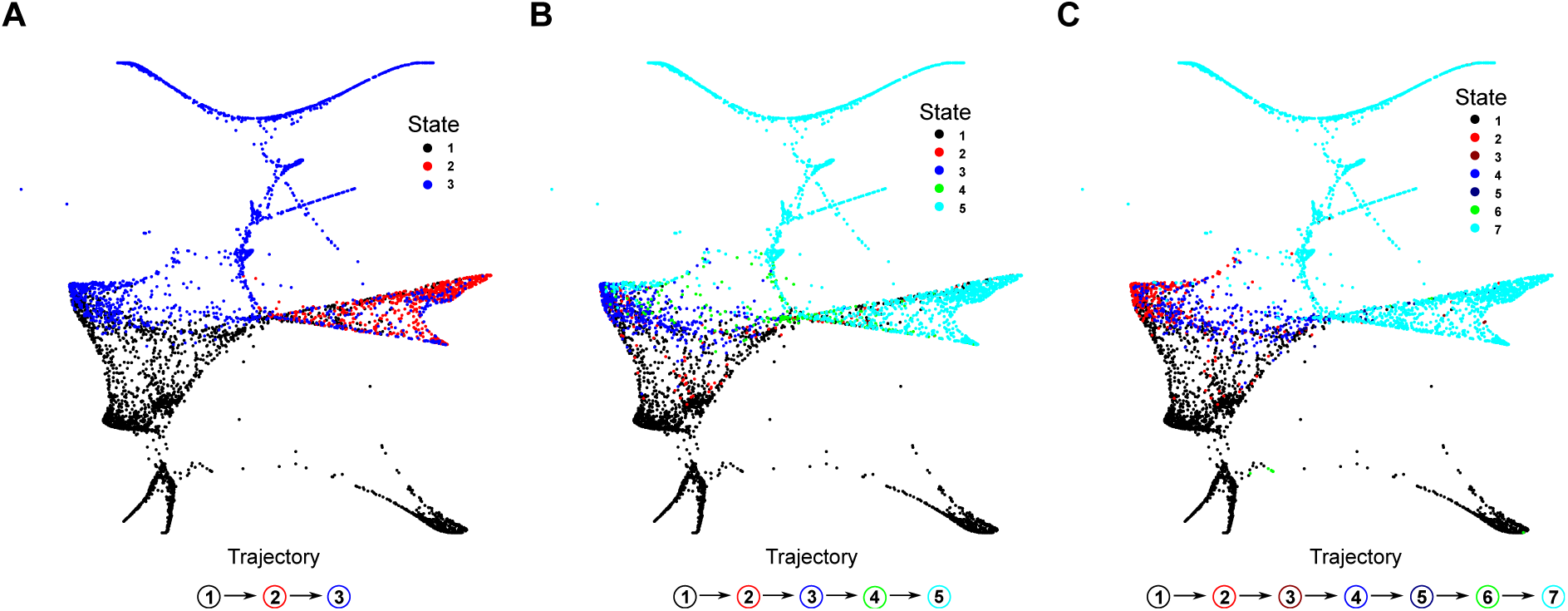
TASIC was run for (A) three, (B) five, and (C) seven states. The results suggest a presence of monolineage trajectories in our data.

### V. Appendix S5

#### Supplemental Notes

This Appendix contains additional notes for analyses in the paper.

##### 1. Definition of gene signatures

From gene set enrichment analysis of 44 gene modules (**table S2**, **fig. S2**), we found significant enrichments for terms that shed light on the reprogramming landscape. We sought to investigate whether we could identify similar expression patterns from well-defined gene signatures. To investigate this, we curated a list of gene sets from various databases of gene signatures (see Table A1, a list of genes for each gene signature is shown in Table S6). A pluripotency gene signature was determined in this work. We performed differential gene expression analysis between two groups of cells: mature iPSCs and cells along the time course D0 to D16 and took the top 100 genes with increased expression in mature iPSCs. A proliferation gene signature was obtained by combining genes expressed at G1/S and G2/M phases. For epithelial and neural gene signatures, we collected canonical markers of epithelial and neuronal cell lineage markers, respectively.

**Table A1.** List of gene signatures used in this work. List of genes for each gene signature are shown in Table S6.

| Gene Signature | Source |
| --- | --- |
| MEF identity | Mouse Gene Atlas (S29, S30) |
| Pluripotency | this work, iPSCs vs. D0 to D16 cells |
| Proliferation | G1/S and G2/M genes, (S31) |
| ER stress | GO:0034976, Biological Process Ontology |
| Epithelial identity | (S32–S35) |
| ECM rearrangement | GO:0030198, Biological Process Ontology |
| Apoptosis | Hallmark P53 Pathway, MSigDB |
| Senescence | Table 1 in (S36) |
| Neural identity | (S37–S43) |
| Placental identity | Mouse Gene Atlas, (S29, S30) |
| X reactivation | chromosome X |

##### 2. Computing descendant distributions for clusters of cells

We compute the descendant distributions for our 33 clusters of cells, some of which span multiple days. To put each cluster on equal footing, we initialize 100 cells in each cluster. We distribute these 100 cells proportionally over the days represented in the cluster.

For each day *d* and cluster *i*, let 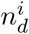 denote the number of day *d* cells in cluster *i*. We denote the total number of cells in cluster *i* by 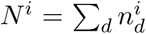. With this notation, we initialize 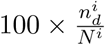 cells in cluster *i* on day *d* and compute the descendant distribution of these cells at the next time point. We denote this descendant distribution by 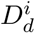. We then compute the mass of this descendant distribution residing in each cluster *j* by summing up the mass 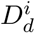 assigns to each cell in cluster *j*. Finally, to obtain the *i, j* entry of the cluster cluster transition table, we sum over *d*.

This give the total mass transferred from from cluster *i* to cluster *j*, per 100 cells initialized in cluster *i*. We compute this separately for 2i and serum.

##### 3. Extraembryonic gene signatures

Previous reports have shown that extraembryonic endoderm stem cells (XEN) were induced in the reprogramming process in parallel of reprograming to iPSCs (S44). To determine if XEN cells were induced in our reprogramming system, we checked the XEN gene signature from in vivo XEN cells, trophoblast and placental gene signatures (**table A2**). While a small fraction of cells (180 cells) displays a high XEN score at day 16 (under serum condition), a larger fraction of cells in clusters 24 and 25 displays high trophoblast and placental signature scores (**fig. A9**). This indicates that the alternative placental-like cell lineage does not share the distinctive XEN signature as previously reported.

**Table A2.** List of XEN, trophoblast and placenta gene signatures.

| Gene Signature | Genes | Reference |
| --- | --- | --- |
| XEN | Dab2 Fst Pdgfra Pth1r Gata6 Foxq1 Fxyd3 Tet3 Sox17 Foxa2<br>Lama1 Lamb1 Gata4 Krt8 | (S45) |
| Trophoblast | Ascl2 Bmp4 Bmp8b Cdx2 Elf5 Eomes Esrrb Ets2 Fgfr2 Grn<br>Igf2 Jade1 Lipg Pcsk6 Ptpra Smad3 Snai1 Tead4 Tfap2c Vav1<br>Yap1 Gata3 Krt7 Krt18 | (S46) |
| Placenta | Table A1 |  |

##### 4. Identifying markers for reprogramming success

To gain further insights into the mechanisms of reprogramming success, we sought to characterize categories of genes that changed their expression in characteristic patterns (**Fig. 5**) along the successful trajectory determined by optimal transport. Genes that exhibited significant changes along the trajectory (2,872 genes) were clustered using *k*-means clustering and the number of clusters was determined by the gap statistic (S47). We identified 14 distinct expression patterns among cells that would end up succesfully reprogrammed (Table S5). Genes were divided into two obvious patterns, upregulated (A1 to A10) and downregulated (A11 to A14). After dox induction, a large number of genes that were mainly involved with MEF identify were downregulated. Instead of “two waves” indicated by a previous report (S48), we observed continuous activation patterns after dox induction. In early stage of reprogramming, they were involved with metabolic changes and were targets of *Myc* (A1 to A3). In late stage (A6 and A7) they were associated with activation of pluripotency networks. We found two categories of pluripotency-associated genes. Genes in category A6 gradually upregulated after dox withdrawal, such as *Nanog*, *Sox2*, *Dppa3* (early pluripotency-associated genes). Genes in category A7 upregulated after genes in A6, such as *Obox6*, *Dppa4* (late pluripotency-associated genes).

**Figure A9.**
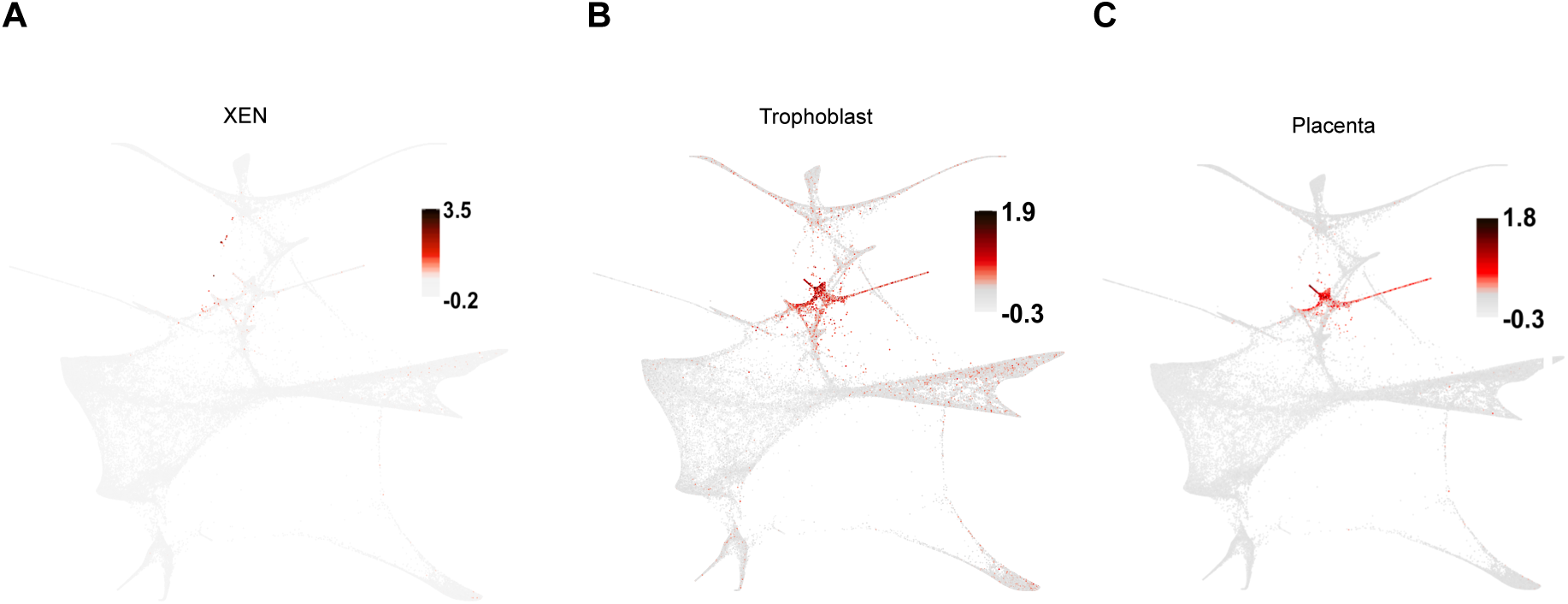
Visualization of single-cell scores for (A) XEN, (B) Trophoblast, and (C) Placenta gene signatures. The color gradient represents the distribution of z-scores across all cells for a given signature.

We next sought to identify genes that were upregulated preferentially in cells that were successfully reprogrammed from A6 and A7. We calculated the fraction of cells in clusters 28 to 33 vs. all other clusters. By setting a threshold of 1%, we ranked genes that were expressed in less than 1% of cells in all other clusters. We found 47 genes that were preferentially expressed in the late stage of reprogramming on successful trajectory and were mostly absent from other cells (Table S5). Future studies could test these candidates as markers for enrichment of fully reprogrammed cells.

##### 5. Cell-cell interactions

To characterize potential cell-cell interactions between contemporaneous cells during reprogramming, we first collected a list of ligands and receptors found in the GO database. The set of ligands (415 genes) is a union of three gene sets from the following GO terms: 1) cytokine activity (GO:0005125), 2) growth factor activity (GO:0008083), and 3) hormone activity (GO:0005179). The set of receptors (2335 genes) is defined by the GO term receptor activity (GO:0004872). Next, we used a curated database of mouse protein-protein interactions (S49) and identified 580 potential ligand-receptor pairs.

We focused on two aspects of potential cell-cell interactions in our data: 1) determining global trends in the expression of all potential contemporaneous ligand-receptors pairs across the reprogramming time course and 2) ranking individual ligand-receptor pairs at a specific day and condition. First, we defined an interaction score *I*_*A,B,X,Y,t*_ as the product of (1) the fraction of cells (*F*_*A,X,t*_) in cluster A expressing ligand X at time *t* and (2) the fraction of cells (*F*_*B,Y,t*_) in cluster B expressing the cognate receptor Y at time *t*. We defined aggregate interaction score *I*_*A,B,t*_ as a sum of the individual interaction scores across all pairs:

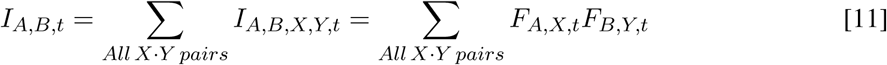

We depicted the aggregate interaction scores for all combinations of cell clusters in **fig. S8A-B**. Second, we sought to explore individual ligand-receptor pairs at a given day and condition between cell subsets of interest. Values of the interaction scores *I*_*A,B,X,Y,t*_ are high for ubiquitously expressed ligands and receptors at a given day and may be nonspecific to a pair of cell subsets of interest. Thus, we used permutations to generate an empirical null distribution of interaction scores between two random groups of cells. In each of the 10,000 permutations, we randomly generated two groups *R*_1_ and *R*_2_ of 100 cells each from time *t* and calculated the interaction score between the ligand in group *R*_1_ and the receptor in group *R*_2_. We then standardized each ligand-receptor interaction score by taking the distance between the interaction score *I*_*A,B,X,Y,t*_ and the mean interaction score in units of standard deviations from the permuted data 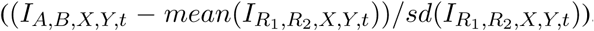. We depicted examples of standardized interaction scores ranked by their values in **fig. S8D-F**.

##### 6. X-chromosome reactivation

We sought to identify X-chromosome reactivation from our scRNA-seq dataset. The set of all detected genes (16,339) was split to X-chromosomal and autosomal genes. Then we calculated the mean X/autosome expression ratio for each cell (normalized by the average X/autosome expression ratio at day 0 cells) as a measurement of X-chromosome reactivation.

The mean X/Autosome expression ratio reached mean value of 1.6 in late stage of reprogramming indicating X-chromosome reactivation (**fig. A10A**). Interestingly, cells in cluster 32 (mature iPSCs in serum) had their X-chromosome inactivated but no *Xist* expression, which might be due to partial differentiation of iPSCs in serum condition or that the established female iPSCs lost one of their X chromosomes, which happens frequently in serum cultured female ESCs or iPSCs but less often in 2i cultured female ESCs/iPSCs (S50). This was specific to mature iPSCs in serum as day-16 cells in serum exhibited similar X-chromosome reactivation to day 16 cells in 2i (**fig. A10B,C**).

Downregulation of *Xist* expression (cluster 28, day 12 cells) preceded X-chromosome reactivation (clusters 29,30,31,and 33; day 16, mature iPSCs) (**fig. A10A-C**). The upregulation of early and late pluripotency genes (activation pattern A6 and A7, respectively) preceded X-chromosome reactivation (**fig. A10D-F**).

We next focused on analyzing the fraction of cells that activated late pluripotency genes A7 and reactivated the X-chromosome. The X/Autosome expression ratio and A7 gene signature score show bimodal distribution across all cells (**fig. A10G** and **fig. A10H**, respectively). We classified cells to those that had reactivated their X-chromosome if the X/Autosome expression ratio *>* 1.4 and those that induced A7 genes if the A7 average z-score *>* 0.25 (**fig. A10G,H**). Using the above thresholds we calculated the fraction of cells in clusters 28-33 that reactivated their X-chromosome and activated the A7 program (**table A3**). Around a 10-fold difference is observed in the percentage of cells that upregulated A7 genes and reactivated X chromosome in clusters 28 and 32.

**Table A3.** Percentages of cells in clusters 28-33 that exhibited X-chromosome reactivation and induction of A7 genes.

| Cluster | 28 | 29 | 30 | 31 | 32 | 33 |
| --- | --- | --- | --- | --- | --- | --- |
| X/A | 7.6 | 79.3 | 84.2 | 89.1 | 7.2 | 81.9 |
| A7 | 72.9 | 98.9 | 99.7 | 99.1 | 93.3 | 99.1 |

##### 7. Identifying large chromosomal aberrations

We have previously developed methods to identify copy number variations (CNVs) in scRNA-Seq data from tumor samples (46). That analysis differed from our current study in two key aspects: (1) the data were higher quality, and sequenced to greater depth in each cell, and (2) we could rely on the clonal expansion of CNVs to make it easier to identify recurring chromosomal aberrations. Because our data are noisy and we don’t expect clonal expansion, we adapted our original methods to carefully control for the statistical significance of aberrations detected in each cell, and to look for non-recurring events.

**Figure A10.**
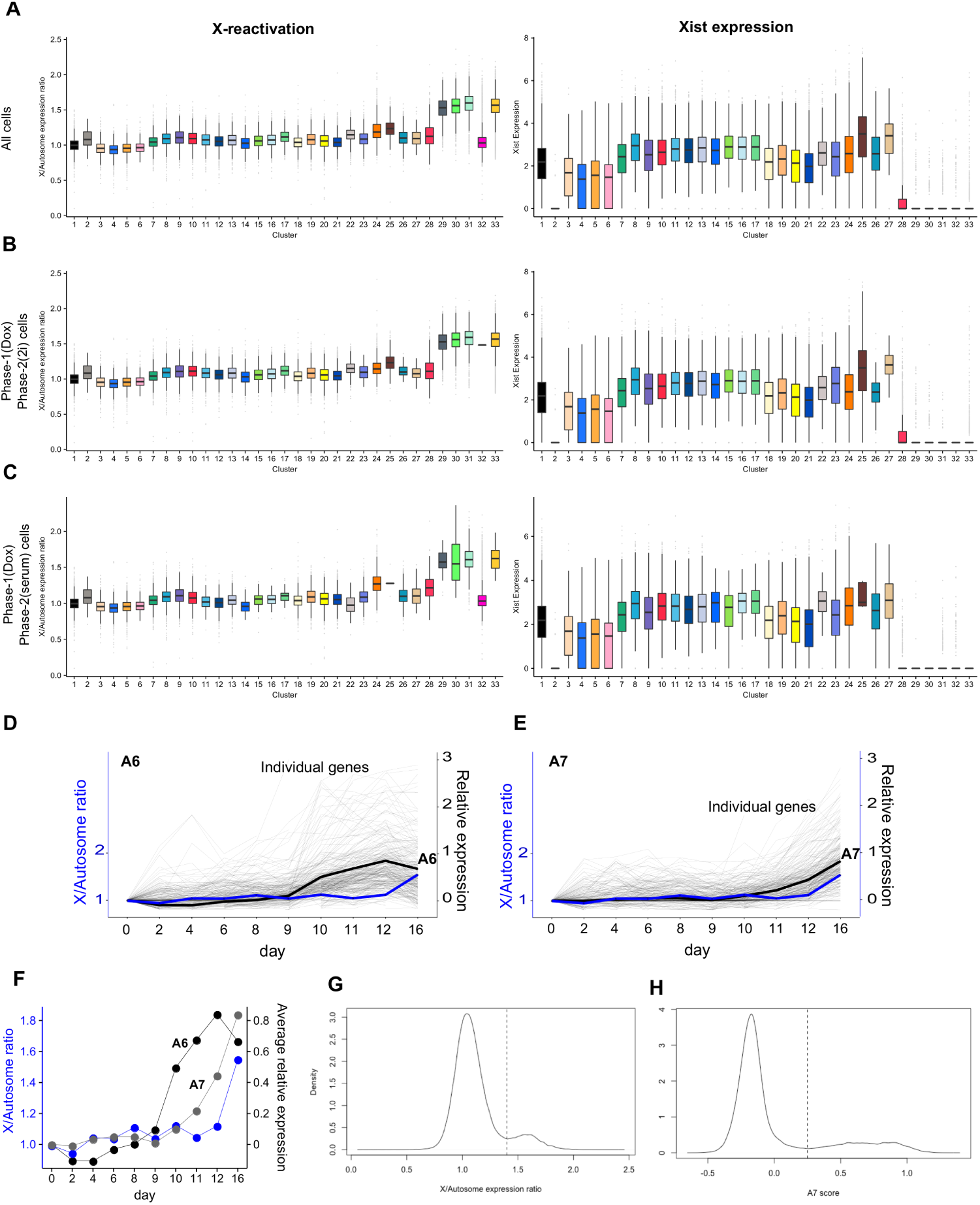
X-chromosome reactivation. (A-C) Boxplots showing X/Autosome expression ratio (left panel) and Xist expression log^2^(E+ 1) across individual cells by clusters (right panel): (A) all cells, (B) phase-1(Dox) and phase-2(2i) cells, (C) phase-1(Dox) and phase-2(serum) cells. (D-F) X/Autosome expression ratio and A6, A7 activation pattern changes along the successful trajectory determined by optimal transport: Relative gene expression changes of individual genes from A6 (D) and A7 (E) activation patterns (gray solid lines). Black and blue solid lines correspond to average relative expression of genes and average X/Autosome expression ratios, respectively. (F) Comparison between activation of A6 and A7 programs (average relative expression) with X/Autosome expression ratio. Distribution of X/Autosome expression ratios (G) and A7 scores (H) across all cells. Dotted lines represent threshold values used in classification of cells that reactivated X-chromosome (*>* 1.4) and upregulated A7 genes (*>* 0.25).

###### Methodology

We performed two types of analysis to detect aberrant expression in large chromosomal regions. First, we searched cells with significant up- or down-regulation at the level of entire chromosomes. Second, we searched for cells with significant subchromosomal aberrations spanning windows of 25 consecutive broadly-expressed genes. Empirical *p*-values and false discovery rates (FDRs) for both analyses were computed by randomly permuting the arrangement of genes in the genome, as described below. Because the X chromosome reactivation is discussed above, we focus on autosomes here.

Permutations for both types of analysis are done as follows. In each of 100,000 permutations we randomly shuffle the labels of genes in the entire dataset, while preserving the genomic positions of genes (with each position having a new label each time) and the expression levels in each cell (so that each cell has the same expression values, but with new labels). We then compute either whole-chromosome or subchromosomal aberration scores for each cell.

To identify whole-chromosome aberrations scores in each cell, we begin by calculating the sum of expression levels in 25Mbp sliding windows along each chromosome, with each window sliding 1Mbp so that it overlaps the previous window by 24Mbp. For each window in each cell, we then calculate the Z-score of the net expression, relative to the same window in all other cells. We then count the fraction of windows on each chromosome with an absolute value Z-score *>* 2. This fraction serves as the whole-chromosome aberration score for each chromosome in each cell. To assign a *p*-value to the whole-chromosome score for celli chromosomej, we calculate the empirical probability that the score for cell_*i*_ chromosomej in the randomly permuted data was at least as large as the score in the original data.

Subchromosomal aberration scores were computed as follows. We begin by identifying the 20% of genes with the most uniform expression across the entire dataset. This is done by calculating the Shannon Diversity (*e*^*entropy*(*gene*)^) for each gene, and taking the 20% of genes with the largest values. Using these genes, we then compute the sum of expression in sliding windows of 25 consecutive genes, with each window sliding by one gene and overlapping the previous window (on the same chromosome) by 24 genes. In each window, we calculate the Z-score relative to all cells at day 0. The net subchromosomal aberration score for a cell is calculated as the l2-norm of the Z-scores across all windows. To assign a p-value to the subchromosomal aberration score for cell_*i*_, we calculate the empirical probability that the score for cell_*i*_ in the randomly permuted data was at least as large as the score in the original data.

For subchromosomal aberration scores we also enriched for chromosomal aberrations (vs. locally coordinated programs of gene expression) by excluding recurrent events. We identified recurrent events by clustering cells based on their aberration profiles (net expression levels across all windows). Clustering was completed by calculating the SVD of all aberration profiles, and performing KMeans clustering on the the top 10 singular vectors (with *k*=100). For each cluster, we quantified cluster compactness and separation using the silhouette score. Cells that were in compact, well-separated clusters (with a silhouette score *>* 0.08) were removed from consideration for subchromosomal aberrations.

For both types of scores, we used the p-values to calculate false discovery rates (FDRs). To identify cells with aberrations at an FDR of *q*, we find the largest p-value, 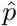, such that 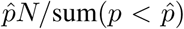, where *N* represents the total number of p-values for a score and sum 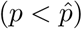 represents the number of p-values less than 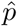.

We applied our method for finding subchromosomal aberrations to an existing single-cell study of oligodendroglioma (S31). Since recurrent aberrations are expected in this setting (due to clonal expansion) we did not remove cells based on clustering recurrent patterns. Applied to these data, our method detected aberrations in 35% of malignant cells (classified in the original study as containing significant copy number variation) and 0% of non-malignant cells (FDR 5%). This demonstrates the specificity and conservative nature of our approach.

###### Results

The results of this analysis are displayed in A11. As we report in the main text, we first looked for whole chromosome aberrations, and found that 0.9% of cells showed significant upor downregulation across an entire chromosome; the expression-level changes were largely consistent with gain or loss of a single chromosome (A11A). Next, when we looked for evidence of large subchromosomal events, we found significant events in 0.8% of cells. The frequency was highest (2.8%) in cluster 14, consisting of cells in the Valley of Stress enriched for a DNA damage-induced apoptosis signature. The frequency was 2-to-3-fold lower in other cells in the Valley (enriched for senescence but not apoptosis), in cells en route to the Valley (clusters 8 and 11), and in fibroblast-like cells at days 0 and 2. Notably, it was much lower (6-fold) in cells on the trajectory to successful reprogramming (A11B,C). Direct experimental evidence would be needed to confirm these events, and to clarify if the aberrations were preexisting in the MEF population, or if they accumulated during the course of reprogramming.

**Figure A11.**
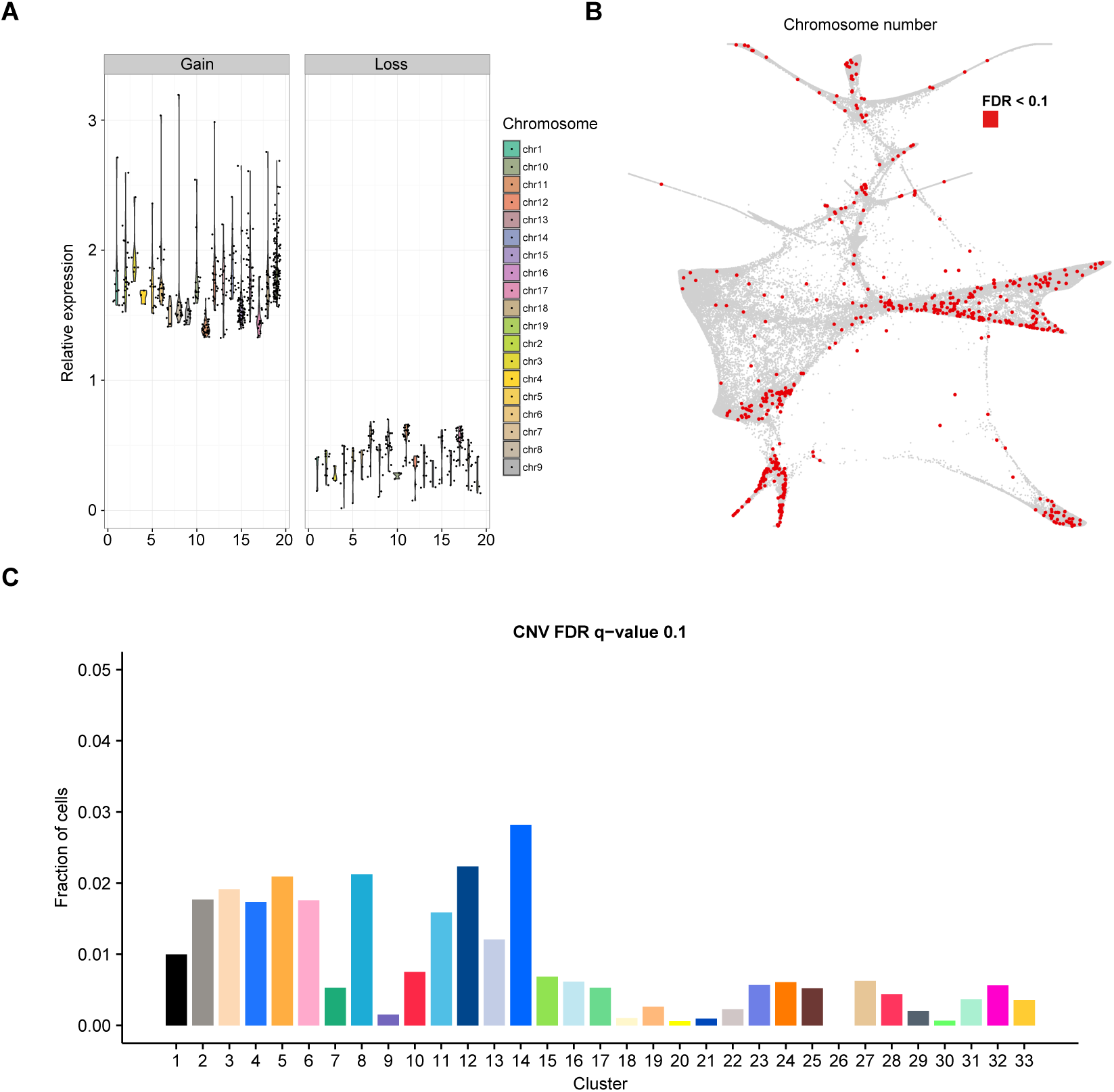
Single-cell expression levels were used to identify cells with aberrant expression in large chromosomal regions. (**A**) Whole chromosome aberrations were detected in 1% of all cells. Each dot represents one chromosome (X axis) in a single cell with significant aberrations (FDR 10%), with violin plots capturing the distributions of dots. The net expression of these chromosomes relative to the average expression across all cells (Y axis) is 1.7-fold higher (median, left panel) and 2.2-fold lower (right panel), indicating whole chromosome gain and loss, respectively. The median relative expression levels are slightly higher (lower) than the 1.5-fold (2-fold) increase (decrease) that would be expected from a true chromosomal gain (loss) because our statistics are conservative in calling significant events but allow for a long tail of high (low) expression. (**B**) Visualization of cells with significant subchromosomal aberrations (red) in FLE. (**C**) Bar plots depict the fraction of cells in each cluster with significant subchromosomal (25-200Mbp) aberrations (FDR 10%).

## VI. Supplementary Figures

Figure S1: **Single-cell RNA-Seq quality metrics.** (**A**) Correlation between number of genes and transcripts per cell (log_10_ transformed). Cells with fewer than 1000 genes detected were filtered out. The color gradient represents cell density. (**B**) Variation in single cell data depicted by correlation between transcript levels (log_10_ transformed average transcript counts) detected in biological replicates generated from day 10 samples in 2i conditions. Pearson correlation coefficient (*r*) is given. The color gradient represents cell density. (**C**) Biological variation in single cell data depicted by correlation between transcript levels (log_10_ transformed average transcript counts) detected in iPSCs and MEFs. Pearson correlation coefficient (*r*) is given. The color gradient represents cell density. (**D**) Correlogram visualizing correlation between single cell gene expression profiles between various time points and their biological replicates. In this plot, the correlation coefficients (circles) are colored according to their values, ranging from 0.75 (blue) to 1 (red). The size of the circles represents the magnitude of the coefficient. The replicates within the timepoints are denoted with suffixes 1 and 2.

Figure S2: **Comparison of various dimensionality reduction methods to visualize single cell RNA-Seq data.** High-dimensional structure of single-cell expression data was embedded in low-dimensional space for visualization using (**A**) the Force-directed Layout Embedding algorithm (FLE) (directed graph approach) and the t-Distributed Stochastic Neighbor Embedding algorithm (t-SNE) with (**B**) principal components and (**C**) diffusion maps as input parameters. See *Appendix 5* for details.

Figure S3: **Visualization of gene modules across reprogramming time points.** Expression profiles of all 65,781 cells studied were embedded in two-dimensional space, using force-directed layout embedding (FLE). The layouts were annotated by single-cell z-scores for 44 gene modules (details in *Table S2*). The color gradient represents the distribution of z-scores across all cells for a given gene module.

Figure S4: **Characterization of cell clusters.** (**A**) Heatmap representing the enrichment of cells from the indicated samples at various time points and culture conditions across 33 different clusters. The color gradient represents the range of cell fractions from 0-0.25. (**B**) Heatmap depicting the enrichment of correlated gene modules within specific cell clusters. The color gradient represents the average gene module scores at the indicated cell clusters. Specific cell clusters that show highly correlated gene module scores were numerically labeled as shown (see *Table S2* for details).

Figure S5: **Visualization of individual gene expression levels.**Normalized expression levels [log_2_(E+1)] for indicated genes were used to annotate force-directed layout embedding (FLE) graphs generated from the expression profiles of 65,781 cells. E represents the number of transcripts of a gene per 10,000 total transcripts.

Figure S6: **Distribution of gene signatures.** (**A**) Distribution of proliferation scores for cells at day 0 (solid black). Proliferation scores were calculated from combined expression levels of G1/S and G2/M cell cycle genes (see *Appendix 5*). Normal mixture modeling (dashed line) was used to classify the cells based on proliferation scores into non-cycling (red) and cycling (blue) cells (top). Visualization of the cycling and non-cycling of cells on FLE at day 0 (bottom). (**B**) Violin plots of single-cell scores for indicated gene signatures and Shisa8 expression levels in clusters 3, 4, 5, and 6. (**C**) Violin plots of single cell scores for indicated gene signatures in clusters 7, 8, and 18. (**D**) Bar plots of normalized expression levels [log_2_(E+1)] for indicated genes, where E is the number of transcripts of a gene per 10,000 total transcripts. (**E**) Single-cell scores for indicated gene signatures across all 33 cell clusters.

Figure S7: **Heatmap depiction of origins and fates of cells inferred from optimal transport.**

Heatmap depiction of cluster descendants in (**A**) serum condition, and cluster ancestors in (**B**) 2i and serum conditions. Each row of the heatmap in (**A**) shows how the descendants of the cells in a particular cluster are distributed over all clusters. Color intensity indicates the number of descendant cells (“mass”, normalized to a starting population of 100 cells) transported to each cluster at the next time point. Each column of the heatmaps in (**B**, **C**) shows how the ancestors of a particular cluster are distributed over all clusters. *Table S5* contains the specific numerical values.

Figure S8: **Potential cell-cell interactions across the reprogramming time course.** (**A**) Temporal pattern of the net potential for paracrine signaling between contemporaneous cells. Each dot represents the aggregated interaction score across all ligand-receptor pairs for a given combination of clusters (all 149 detected ligands). The aggregate interaction score is defined as a sum of individual interaction scores. (**B**) As in **A**, but genes specific to SASP signature are considered (20 detected ligands). (**C**) Heatmap representing the aggregate interaction scores on day 16 cells in 2i condition for ligands specific to SASP signature. Rows correspond to clusters of cells expressing ligands. Columns correspond to clusters of cells expressing cognate receptors. Only clusters containing more than 1% of cells from day 16 (2i) are shown. (**D**-**F**) Potential ligand-receptor pairs ranked by their standardized interaction scores calculated from the permuted data (see *Appendix 5* for details). Ligand-receptor pairs between valley of stress cells (clusters 11-17) and iPSCs (clusters 28-33) on day 16 (2i), (**E**) valley of stress cells and preneural/neural-like cells (clusters 23, 26, and 27) on day 16 (serum), and (**F**) placental-like cells (clusters 24 and 25) and valley of stress cells on day 12 (2i).

Figure S9: **Gene modules and associated transcription factors based on optimal transport.** Using optimal transport trajectories, TF levels in cells at time *t* are used to predict the activity levels of gene modules in descendant cells at time *t* + 1. Gene modules are learned during model training to capture coherent expression programs. For five modules (**A**-**E**), bar plots depict the top 50 genes in the module (black), and the top 20 TFs each associated with positive (red) and negative (blue) module activity. (**AB**) Two modules that are active in cells with placental identity. (**C**) A module active in cells with neural identity. (**D**-**E**) Two modules active in successfully reprogrammed cells. (**F**) Enrichment analysis of TFs in day 12 cells with high (*>*80%) vs. low (*<*20%) probability of successful reprogramming. Dot size and color represent percentage of day 12 cells expressing the indicated TF in high- or low-probability cells. Bar heights indicate the fold enrichment in highvs. low-probability cells.

Figure S10: **Effect of overexpression of *Obox6* and *Zpf42* on reprogramming efficiency.** (**A**) Percentage of Oct4-EGFP^+^ cells at day 16 of reprogramming from secondary MEFs by lentiviral overexpression of *Oct4*, *Klf4*, *Sox2*, and *Myc* (OKSM) combined with either *Zfp42*, *Obox6*, or an empty control, in either 2i or serum conditions. Oct4-EGFP^+^ cells were measured by flow cytometry. Plot includes the percentage of Oct4-EGFP^+^ cells in three biological replicates (for *Zfp42* and *Obox6* overexpression, or an empty control) from five independent experiments (Exp). (**B**, **C**) Number of Oct4-EGFP^+^ colonies at day 16 of reprogramming from primary MEFs by lentiviral overexpression of individual *Oct4*, *Klf4*, *Sox2*, and *Myc* combined with either *Zfp42*, *Obox6*, or an empty control in (**B**) 2i and (**C**) serum conditions. Plot includes the number of Oct4-EGFP^+^ cells in three biological replicates (for *Zfp42* and *Obox6* overexpression, or an empty control) from two independent experiments (Exp).

## VII. Supplementary Tables

Table S1: **Summary of single cell sequencing statistics and sample information.**

Table S2: **List of genes for each module and gene set enrichment analyses for specific gene modules highlighted in the test.**

Table S3: **List of the percentage of cells in each cluster at different collection time points.**

Table S4: **List of genes comprising gene signatures indicated in Fig. 3E.**

Table S5: **Descendant and ancestral distributions for clusters, in 2i and serum conditions.**

Table S6: **List of genes and gene set enrichment analyses for 14 activation groups along the successfully reprogrammed trajectory reported in Fig. 5.**

Table S7: **TFs enriched in cells with a high probability of iPSC, neural, or placental fate.**

## VIII. Supplementary Videos

S Video1: **Video highlighting cells in 2i at specific collection time points during the reprogramming process on force-directed layout embedded maps.**

S Video2: **Video highlighting cells in serum at specific collection time points during the reprogramming process on force-directed layout embedded maps.**

## Footnotes

1 A limitation of Waddington’s landscape is that it is cell-autonomous (i.e. doesn’t include effects of other cells). Our model is actually more general.

2 For example, one simple cost function is squared Euclidean distance *c*(*x, y*) = ||*x - y*||^2^.

3 For example, imagine that ℙ*t* is a mixture of Gaussians with time-varying mixture weights.

4 *Advection*, a term borrowed from fluid mechanics, refers to the transport of a substance by bulk motion. The constraint that the divergence of the flow is equal to the rate of change of *ρ* means that *ρ* flows according to the velocity field *v*, without gaining or losing mass.

5 As we discuss in Appendix S2, we have had some initial success with a rectified-linear function class. Following Cleary et al. 2017 [SMAF], we impose low-rank and sparsity constraints on the linear part, and we apply the function only to the transcription factors in *Xti*. Each low-rank component can then be interpreted as a regulatory module of transcription factors acting on a module of regulated genes.

